# A recipient-based anti-conjugation factor triggers an abortive mechanism by targeting the Type IV secretion system

**DOI:** 10.64898/2026.03.10.710775

**Authors:** Liyana Ayub Ow Yong, Jiang Yeow, Suma Tiruvayipati, Siyi Chen, Celestine Grace Xueting Cai, Swaine Lin Chen, Shu-Sin Chng

**Affiliations:** Infectious Diseases Translational Research Programme, National University of Singapore, Singapore; Department of Medicine, Division of Infectious Diseases, Yong Loo Lin School of Medicine, National University of Singapore, Singapore 119074; Laboratory of Bacterial Genomics, Genome Institute of Singapore, Agency of Science, Technology and Research (A*STAR), Singapore 138672; Department of Chemistry, Faculty of Science, National University of Singapore, Singapore 117543; Biomolecular Sequence to Function Division (BSFD), Bioinformatics Institute (BII), Agency for Science, Technology and Research (A*STAR), Singapore, Singapore 138671; Singapore Centre for Environmental Life Sciences Engineering, National University of Singapore (SCELSE-NUS), Singapore 117456

## Abstract

Many bacterial defense (immune) systems prevent the entry of foreign DNA by directly recognizing and targeting nucleic acids, effectively blocking all mechanisms of horizontal gene transfer^1^. However, systems defending specifically against conjugation, a major route for gene dissemination, have heretofore not been reported. We have discovered a novel defense factor, which we name AbjA (**<u>Ab</u>**ortive con**<u>j</u>**ugation protein **<u>A)</u>**, that specifically limits successful plasmid conjugation into a recipient bacterium. AbjA interacts directly with and targets the ATPase component TrbE of the Type IV secretion system (T4SS)^2,3^ to induce cell death; this contrasts with most other defense systems that act at the nucleic acid level. AbjA therefore represents the first member of a new class of bacterial defense factors that trigger what we term ‘abortive conjugation’. Previously, recipient bacteria were viewed largely as defenseless against the mechanism of conjugation^4–6^; our discovery and characterization of AbjA demonstrates that recipient bacteria can block conjugation to limit the transfer (and thus spread) of plasmids. Discovery of this class of defense systems thus has implications for bacterial defense, plasmid evolution, and possible strategic alternatives to rationally target plasmid spread, particularly with respect to virulence and antibiotic resistance.

## Introduction

Conjugative plasmids are autonomous genetic entities that can facilitate the rapid dissemination of virulence factors and antibiotic resistance genes (ARGs) across bacterial populations. Conjugation is mediated by a class of specialized transfer apparatus called the Type IV secretion system (T4SS), which comprises 11 core proteins (VirB1-VirB11, as designated in T4SS archetype *Agrobacterium tumefaciens*)^7^ that assemble into an envelope-spanning substrate translocation channel^8^ and a dynamic surface appendage called the pilus, which extends and retracts to establish contact with potential recipient cells^9–11^. Pilus contact eventually results in formation of a mating pair, after which the plasmid enters the recipient cell, establishes itself, and enables the cell to participate in further rounds of horizontal gene transfer (HGT) by expressing the T4SS anew^12^. The energy for DNA transfer is supplied by three conserved ATPases: VirD4, VirB11, and VirB4. Of these, VirB4 is present in all of the Gram-negative T4SSs found in conjugative plasmids and is therefore a useful marker of plasmid transmissibility and a reasonable proxy for evolutionary relatedness^13^.

Conjugation is one of four main mechanisms by which bacteria can acquire new (foreign) DNA^14,15^. Foreign DNA itself may confer adaptive advantages, such as resistance to antimicrobials, but expression and maintenance of new genes are associated with significant metabolic costs, often compromising bacterial fitness in the absence of selective pressure to maintain the incoming DNA^16,17^. Therefore, bacteria have developed a variety of defense mechanisms that can prevent the entry or establishment of foreign DNA. Some of these systems can prevent the transfer of conjugative plasmids, but none act specifically on conjugation and none are known to interfere specifically with the conjugative machinery. Key examples include the recently characterized DdmABCDE and ApsAB systems that reduce conjugation frequency when present in the recipient bacterium; they also prevent non-conjugative HGT mechanisms, such as phage infections and the transformation of small, high-copy number plasmids^18–20^. Other general DNA defense systems such as restriction-modification (RM) may in principle block conjugation, but also other HGT routes^21–24^. Similarly, despite evidence that spacer content can target conjugative plasmids, Type IV CRISPR-Cas systems are manifestly not specific against conjugation, appearing more to be repurposing a general nucleic acid-based system for targeting. Furthermore, most of these systems are not encoded on the chromosome of the recipient bacterium but instead carried on conjugative plasmids themselves, where they are thought to mediate inter-plasmid competition, prioritising plasmid dominance rather than host protection^25,26^.

The above-mentioned defence systems that can target conjugative plasmids are triggered by recognition of specific DNA or RNA sequences. This mode of action overlaps with that of many phage-targeting systems, such as RM, CRISPR-Cas, and prokaryotic argonautes (pAgo)^27^. Upon sequence detection, these systems deploy diverse molecular mechanisms, including abortive infection^28–30^, inhibition of phage replication or transcription^28,31–33^, and nucleic acid cleavage^28,34–36^. Further afield, offensive systems like the Type VI secretion systems may be triggered by conjugation (i.e. defense by counterattacking the donor cell), but again the mechanism of inhibition does not directly target conjugation^37^. Thus, there are as yet no described examples of conjugation-specific defense systems, nor of systems that can inhibit conjugation through a mechanism other than the general one of direct DNA recognition and degradation. Indeed, it is commonly thought that recipient cells are helpless to prevent incoming conjugation^4–6^.

In this study, we identify a recipient-encoded factor that markedly reduced the transfer frequency of specific conjugative plasmids by inducing cell death during plasmid establishment. Akin to conceptually similar mechanisms in phage defense systems, we term this process “abortive conjugation” and name this factor **<u>ab</u>**ortive con**<u>j</u>**ugation protein **<u>A</u>** (AbjA). In recipients encoding *abjA*, the T4SS ATPase, TrbE (a VirB4 homolog), from the incoming plasmid is essential for triggering abortive conjugation. We demonstrate that AbjA directly interacts with TrbE; this interaction disrupts the latter’s oligomerization and function, possibly leading to runaway hydrolytic activity and cell death. AbjA represents the first example of a defense mechanism that is (i) specific against conjugation, and (ii) not based on nucleic acid recognition. As in both anti-phage abortive infection and eukaryotic viral defense systems, AbjA should limit the penetration of targeted conjugative plasmids into the broader recipient population by means of self-sacrifice. The discovery of AbjA further suggests that anti-conjugation targets may represent viable alternatives for limiting the spread of antimicrobial resistance.

## Results

### *abjA* is necessary and sufficient to inhibit plasmid transfer of specific conjugative plasmids

A challenge in discovering naturally-occurring defense systems against conjugation is a lack of a universal donor strain, i.e. one that can conjugate with most recipient strains, yet still enable efficient selection of transconjugants. In contrast, universal recipient strains like *Escherichia coli* J53, having relatively unique resistance to azide, are well known and have enabled discovery of many new and diverse conjugative plasmids. We therefore leveraged an efficient inducible toxin-based negative selection system (P*_rhaB_*-*tse2*)^38^, to convert *E. coli* K12 strain MG1655 into such a universal donor strain. The high efficiency of this particular negative selection system is crucial for enabling conjugation screens using unmarked, wild-type recipients. Furthermore, it achieves this selection efficiency in the context of minimal media, which may better mimic typical environmental niches than rich media (Fig. 1a). Thus, in searching for recipient-based conjugation-modifying factors, we were able to utilize essentially any strain (effectively enabling access to the natural broad diversity of bacteria strains, both at the level of allelic variation and variable gene presence) in alternative media conditions, offering distinct advantages compared to previous studies. We initially performed a simple screen for low efficiency conjugative plasmid transfer against a panel of well-studied, unmodified clinical strains; we found that the uropathogenic *E. coli* strain CFT073 was a poor recipient (by a factor of one million) for the broad-range conjugative plasmid pRK24^39,40^ in low-nutrient conditions (Fig. 1b).

**Figure 1.**
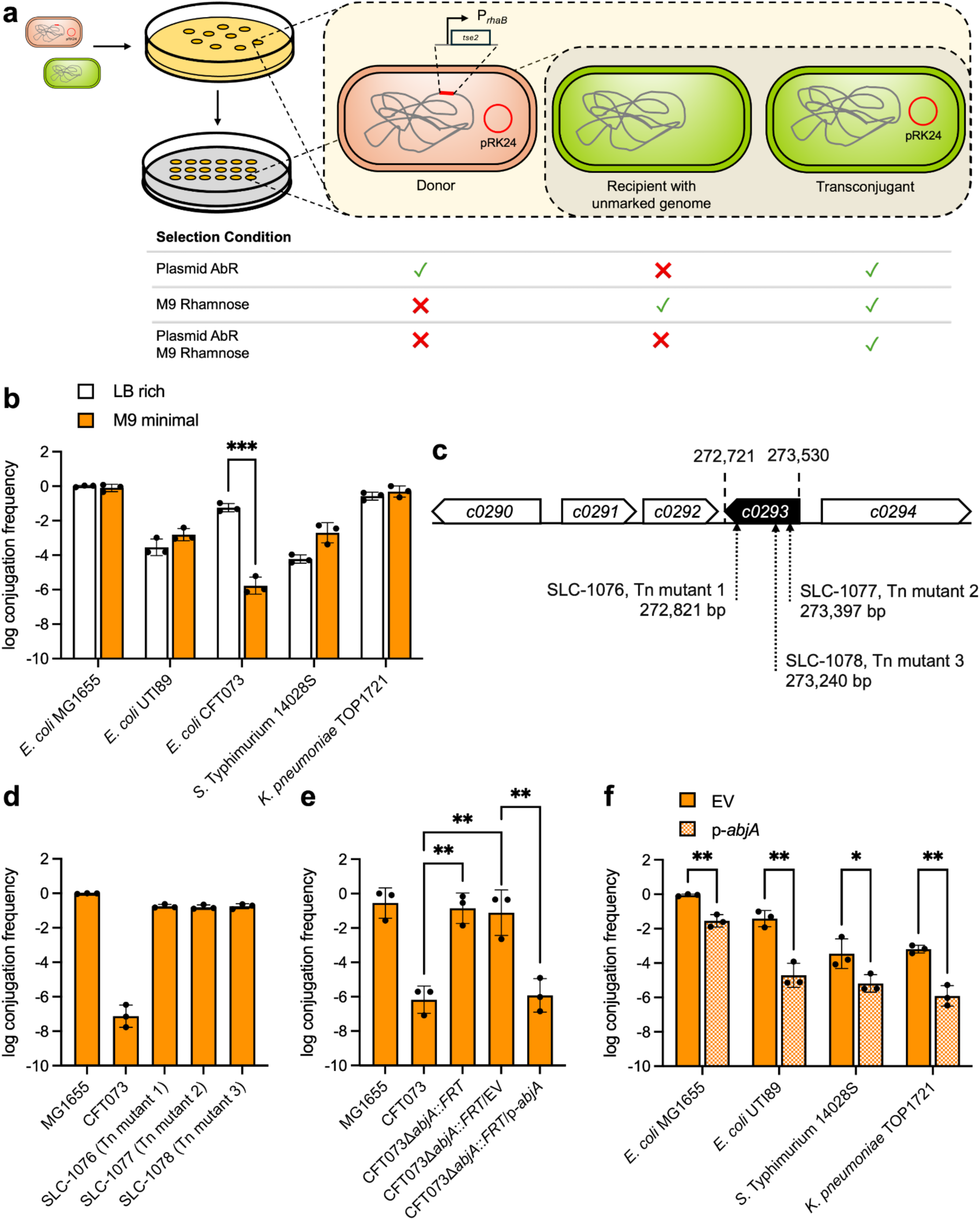
*abjA* is necessary and sufficient to inhibit conjugative transfer of pRK24. (a) Overview of our negative-selection-based conjugation setup. A negative selectable marker (P*rhaB*-*tse2*) was inserted in the chromosome of the donor strain to facilitate transconjugant selection. Upon 1 hour of solid mating, transconjugants were selected for on minimal media supplemented with rhamnose, which eliminates the donor cell population, along with the appropriate antibiotics to remove the recipient cell population. (b) Conjugation frequency of pRK24 from MG1655 into various clinical strain recipients in rich and minimal media. (c) Tn5 insertions that recovered conjugation efficiencies in all three CFT073 clones were found within *c0293* (*abjA*) as indicated. Conjugation frequency of pRK24 from MG1655 into (d) CFT073 Tn5 mutants, (e) CFT073Δ*abjA*, and (f) other strains and species that do not encode *abjA*, harboring pACYC177 empty vector (EV) or pACYC177-*abjA*-*his* (p-*abjA*) as indicated, in minimal media. Log conjugation frequency data are presented as mean ± s.d. of three biological repeats. Unpaired Student’s t-test: *, p <0.05, **, p < 0.01, ***, p < 0.001.

To identify the genetic basis for this lower rate of conjugation, we switched to positive selection, screening a transposon mutant library of CFT073 for mutants with an increase in plasmid transfer efficiency. We discovered that disruption of a single gene, *c0293* (which we hereafter refer to as *abjA* (for **<u>ab</u>**ortive con**<u>j</u>**ugation protein **<u>A</u>**)), improved pRK24 transfer efficiency by ∼six orders of magnitude (Fig. 1c, d). We further verified that a CFT073 strain carrying a targeted deletion of the *abjA* gene was an efficient conjugation recipient, and in this deletion strain, expression of *abjA in trans* restored the low conjugation efficiency phenotype without impacting growth (Fig. 1e, Supplementary Fig. S1). Therefore, *abjA* is necessary for this putative conjugation (of pRK24) inhibition phenotype in CFT073.

We next tested whether *abjA* itself is sufficient to inhibit conjugation. Indeed, in other *E. coli* strains that do not encode *abjA*, heterologous expression substantially reduced the frequency with which successful transconjugants with pRK24 were isolated. (Fig. 1f). The same was true in *Salmonella* enterica Typhimurium 14028S and *Klebsiella pneumoniae* TOP1721. Furthermore, expression of *abjA* in the recipient strain also inhibited the transfer of two other conjugative plasmids, R751 and R388, using MG1655 as both donor and recipient (Fig. 2a). We also assessed whether *abjA* could work against other forms of HGT, e.g. laboratory-induced transformation. Remarkably, we found that *abjA* blocked plasmid transfer during transformation, but only of pRK24, R751, and R388 (Fig. 2b), reproducing similar selectivity. These results demonstrate that the effect of *abjA* is in principle independent of the transfer process, but likely exerted during or after plasmid establishment. We sought to determine if *abjA*-mediated transfer inhibition could be attributed to known recipient-based defense mechanisms. When CFT073 was used as both donor and recipient, we saw similar levels of conjugation inhibition, suggesting that *abjA* is unrelated to any RM system (Supplementary Fig. S5). Moreover, no CRISPR-Cas locus was identified in the CFT073 genome^41^. We thus conclude that *abjA* is necessary and sufficient in blocking conjugative plasmid transfer, presumably via a novel mechanism.

**Figure 2.**
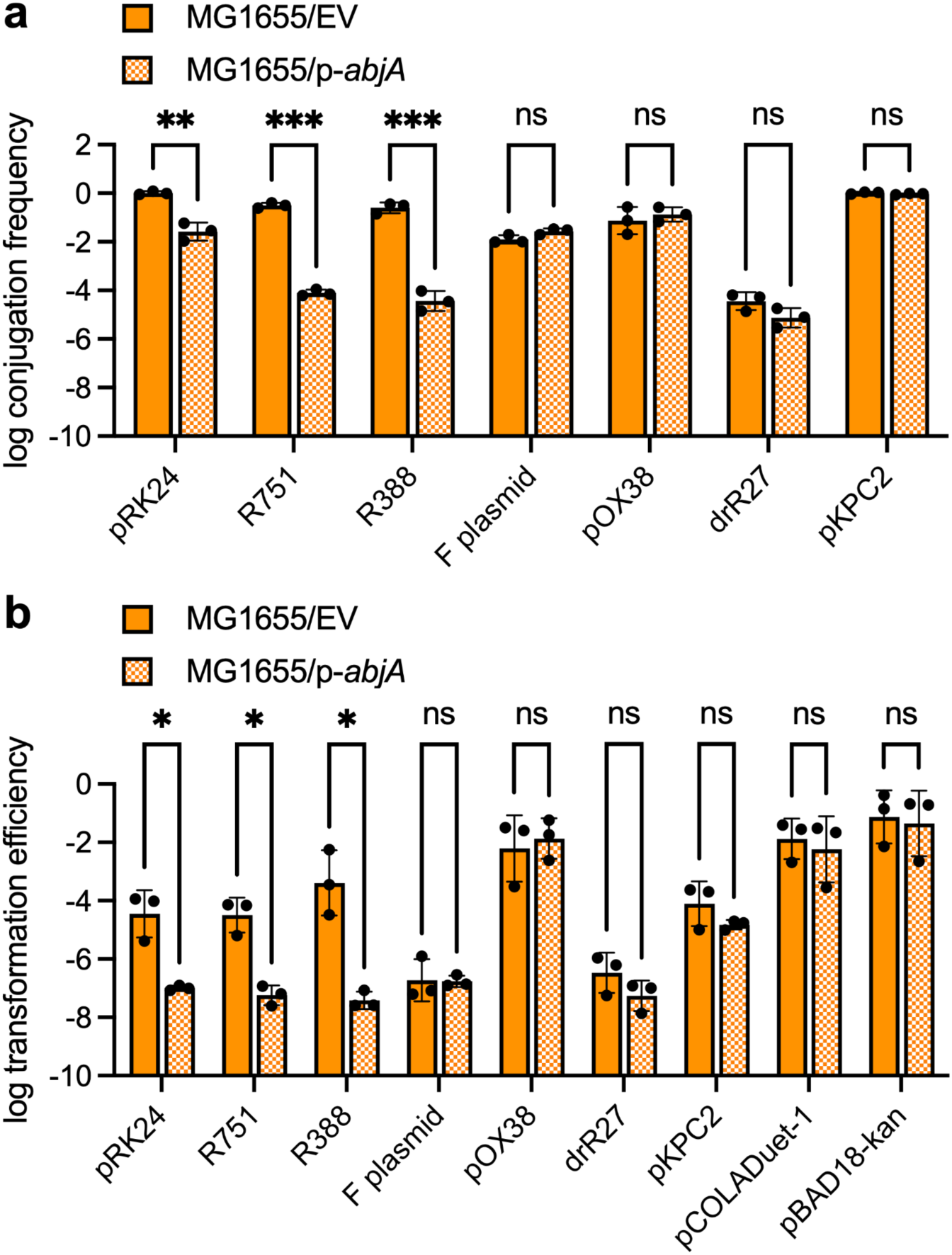
*abjA* inhibits plasmid transfer via conjugation and transformation, selectively affecting only a subset of plasmids. (a) Conjugation frequency (a) and transformation efficiency (b) of various conjugative plasmids (see Supplementary Table S3) into MG1655 recipient harboring pACYC177 empty vector (EV) or pACYC177-*abjA*-*his* (p-*abjA*). Log conjugation frequency and transformation efficiency data are presented as the mean ± s.d. of three biological repeats. Unpaired Student’s t-test: *, p <0.05, **, p < 0.01, ***, p < 0.001, ns, not significant.

Given the conjugation inhibition phenotype, one might expect to find *abjA* or homologs in other bacterial species. Interestingly, we found *abjA* to be rare, occurring in only 0.8% of a pooled dataset of 178,720 genomes and assemblies, and largely restricted to *E. coli* (Supplementary Fig. S2a), particularly sequence type 73 (ST73) (Supplementary Fig. S2b). We also observed that *abjA* frequently co-occurred with three upstream genes, *c0290*, *c0291*, and *c0292* (86/118; Fig. 1c), although these did not contribute to the inhibitory function (Supplementary Fig. S3). Of these, *c0291* encodes a putative integrase, suggesting a possible method for how *abjA* could be chromosomally acquired and/or spread. Outside of *E. coli*, a genetically distinct group of *abjA* homologs can be found in a *Salmonella* cluster (Supplementary Fig. S4a; nucleotide identity ∼91%). We confirmed that these *Salmonella* alleles encoded functional *abjA* defense systems because heterologous expression of these genes in *E. coli* was sufficient to inhibit pRK24 transfer via transformation (Supplementary Fig. S4b). Since functional *abjA* can be found in multiple species of Enterobacteriaceae, it is tempting to speculate that there may be more divergent sets of genes that could be evolutionarily or mechanistically related to *abjA* (as there are multiple classes of RM and CRISPR-Cas systems), that we simply have yet to find.

### Conjugative plasmid establishment in *abjA*-bearing cells induces cell death

We consistently observed that the presence of *abjA* was associated with a 1-2 (and up to 6) log reduction of recipient CFU during conjugative transfer of pRK24 (Supplementary Fig. S6). We therefore hypothesized that the co-occurrence of *abjA* and pRK24 was likely incompatible for cell viability. To test this idea, we placed *abjA* under an inducible promoter to determine its effect on fitness in MG1655 in the presence of pRK24. Consistently, plating efficiency was drastically reduced only when *abjA* expression was induced in the presence of pRK24 (Fig. 3a). In liquid media, these cells exhibited a significant lag phase (∼8 h) and only began exponential growth thereafter, possibly due to genetic suppressors (Fig. 3b) (see below). Cells expressing *abjA* in the presence of pRK24 were highly filamentous, with unsegregated chromosomes, possibly leading to eventual cell death (Fig. 3c). We showed that CFT073 carrying pRK24 also exhibited plating defects and were similarly elongated under low-nutrient conditions, establishing the relevance of these *abjA*-associated phenotypes in the native host (Supplementary Fig. S7a,b).

**Figure 3.**
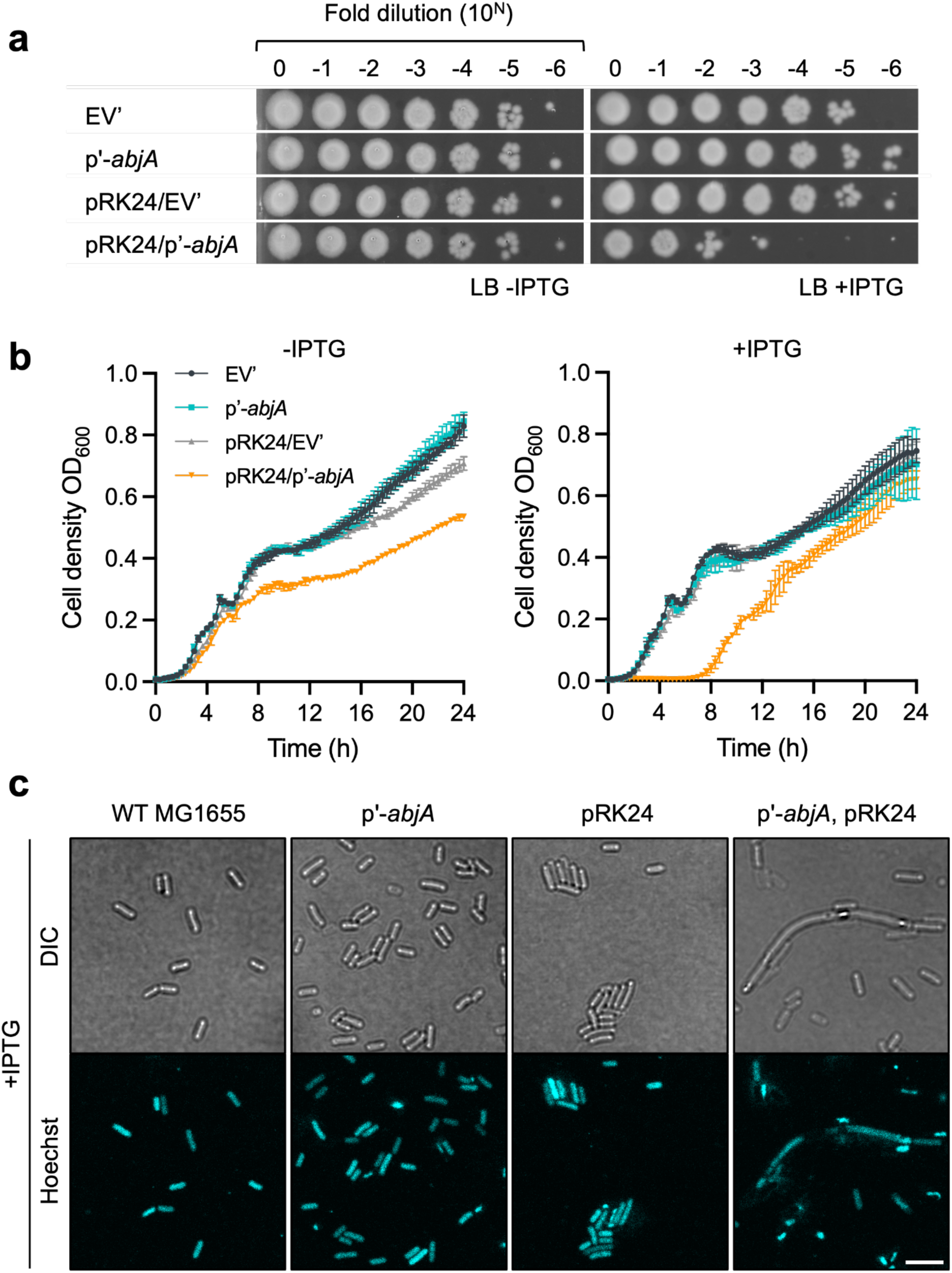
Co-occurrence of pRK24 and *abjA* are incompatible for cell viability. (a) Efficiency of plating (EOP) and (b) growth curves of MG1655 cells harboring pTrc99A (EV’), or pTrc-*abjA*-*his* (p’-*abjA*), with or without pRK24, in LB. 1 mM IPTG was added (as indicated) to induce overexpression of *abjA*. (c) DIC and fluorescence microscopy images of strains from (b) grown in the presence of 1 mM IPTG for 5 hours. Intracellular DNA was labelled with Hoechst stain. Scale bar represents 5 *µ*m.

### The mode of action of AbjA depends on the T4SS factor TrbE

We hypothesized that one or more gene(s) or gene product(s) on pRK24 is incompatible with *abjA*, thus causing cell death during plasmid establishment. To test this idea, we first generated deletions in regions of pRK24 that are known to not impact plasmid maintenance. Since deletions within the transfer gene regions will impact conjugation, we used the transformation assay described above to circumvent conjugation per se and directly assess the resultant plasmid transfer inhibition phenotype. The deletions overall encompassed ∼56% of the ∼60 kb comprising pRK24, including the distinct IncP transfer gene regions Tra1 (Δ*traA-X*) and Tra2 (Δ*trbA-P*) as well as the putative lethal gene regions (Δ*klaABC*, Δ*kleAB*, and Δ*kleCDEF*) (Fig. 4a, Supplementary Table S4). Interestingly, only the transformation of the pRK24Δ*trbA-P* construct was no longer inhibited by the presence of *abjA* (Fig. 4b, Supplementary Fig. S8), confirming that any such critical gene(s) should not be in the Tra1 or lethal gene regions. Subsequent deletions of *trbEFG*, and then single-gene deletions, revealed all genes except *trbE* to be unrelated to preventing transformation of pRK24 in the presence of *abjA* (Fig. 4c). pRK24 constructs that lack *trbE* no longer caused plating defects, nor a long lag growth phase, when *abjA* was overexpressed (Fig. 4d,e). Notably, deleting *trbE* from the other targeted plasmids, R751 and R388, abolished *abjA*-dependent inhibition of transformation, plating defects, and growth delays (Supplementary Fig. S9). Altogether, these results identify *trbE* as the only possible key gene among these regions that determines the fate of the *abjA*-expressing recipients when receiving these plasmids.

**Figure 4.**
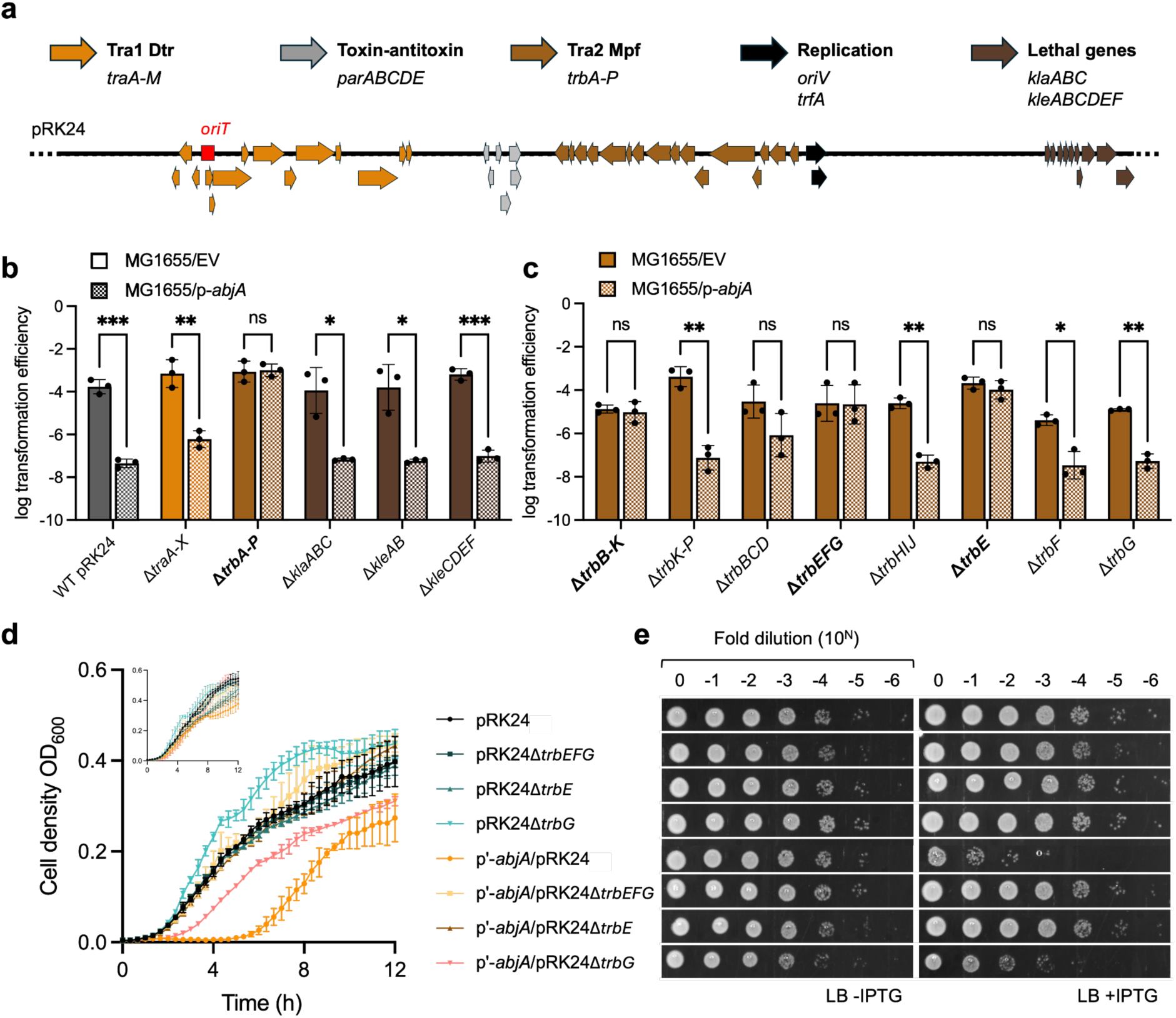
The single gene *trbE* (encoding the ATPase for T4SS) on pRK24 is necessary to cause incompatibility with *abjA*. (a) Schematic of the major features of pRK24 (see Supplementary Table S4 for details). (b) and (c) Transformation efficiencies of pRK24 constructs bearing various different gene deletions into MG1655 recipient harboring pACYC177 empty vector (EV) or pACYC177-*abjA*-*his* (p*-abjA*). Log transformation efficiency data are presented as the mean ± s.d. of three biological repeats. Unpaired Student’s t-test: *, p <0.05, **, p < 0.01, ***, p < 0.001, ns, not significant. (d) Growth curves and (e) EOP of various MG1655 strains harboring pTrc (EV’) or pTrc-*abjA*-*his* (p’-*abjA*) together with indicated pRK24 constructs, with and without the supplementation of 1 mM IPTG on LB.

Beyond the extensive lag phase, cells harboring unmodified pRK24 in the presence of *abjA* eventually entered exponential growth phase (Fig. 3b). This pattern suggests that the cells that eventually grew did so because they had accumulated suppressor mutations. We isolated and sequenced plasmids from ten individual clones after 24 h. No mutations were found within *abjA*. Remarkably, pRK24 from all ten “suppressors” contained only mutations in *trbE*, represented by six unique variants that mostly resulted in C-terminal truncations (Supplementary Table S6). These pRK24 suppressor variants were also no longer inhibited by *abjA* during transformation (Fig. 5a), nor caused plating defects and delayed growth (Fig. 5b,c). Combined with the gene deletion results, we conclude that *trbE* is necessary for the *abjA*-mediated reduction in plasmid transfer efficiency. Finally, we demonstrated that the transformation of an unrelated plasmid expressing *trbE*_pRK24_ alone was effectively inhibited by *abjA* (Fig. 6a), establishing that *trbE* is sufficient to block plasmid transfer.

**Figure 5.**
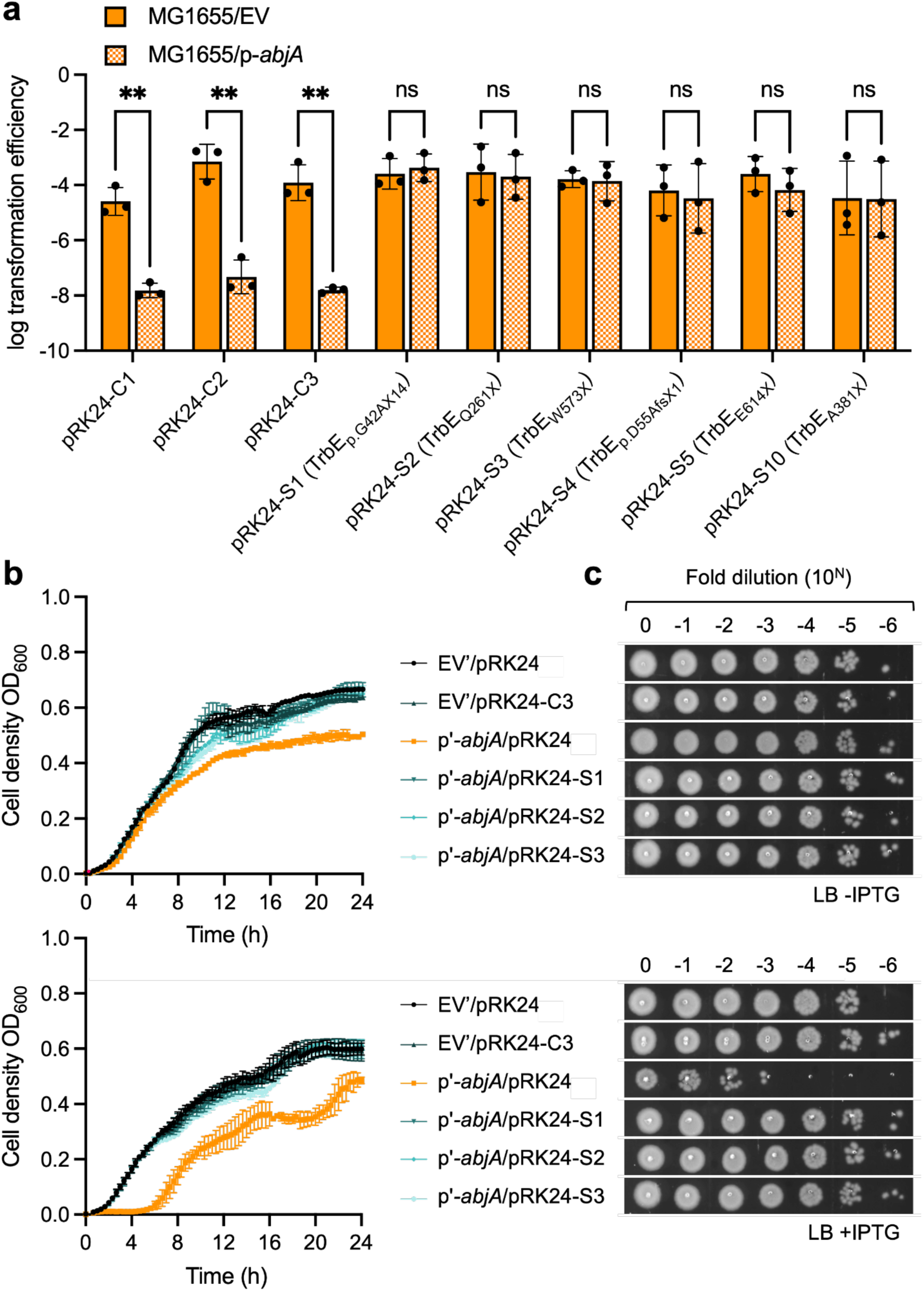
Suppressor mutations within pRK24 *trbE* caused a loss of *abjA-*associated lethal phenotypes. (a) Transformation efficiencies of pRK24 (isolated from overnight cultures in the absence (pRK24-Cx series), or presence (pRK24-Sx series) of *abjA*), into MG1655 recipient harboring pACYC177 empty vector (EV) or pACYC-*abjA*-*his* (p*-abjA*). Log transformation efficiency data are presented as the mean ± s.d. of three biological repeats. Unpaired Student’s t-test: **, p < 0.01, ns, not significant. (b) Growth curves and (c) EOP of various MG1655 strains harboring pTrc (EV’) or pTrc-*abjA*-*his* (p’-*abjA*) together with indicated pRK24 constructs, with and without the supplementation of 1 mM IPTG on LB.

**Figure 6.**
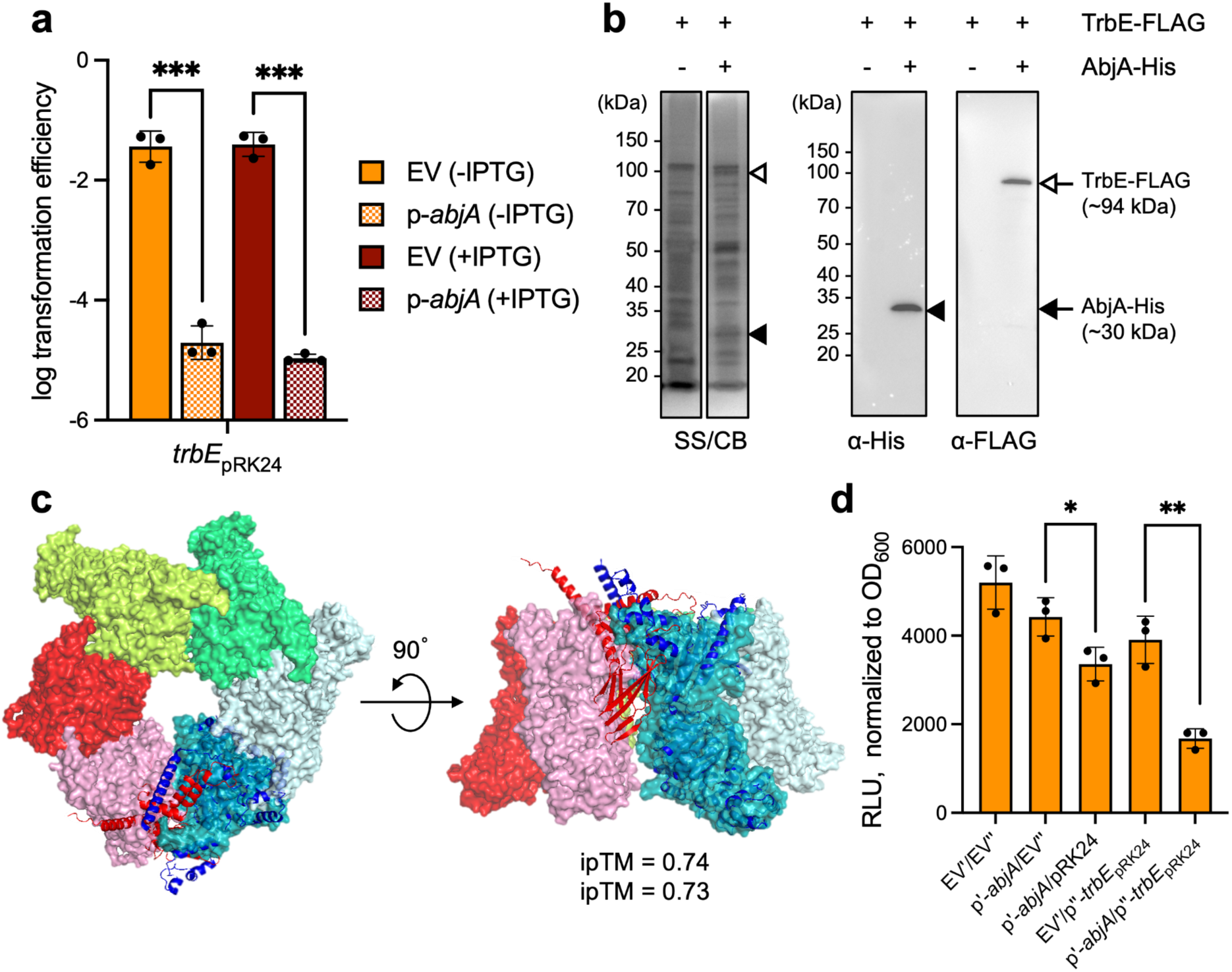
TrbE interacts directly with AbjA to cause cell lethality. (a) Transformation efficiency of pCOLADuet-*trbE*-FLAG (*trbE*_pRK24_) into MG1655 recipient harboring pACYC177 empty vector (EV) or pACYC177-*abjA*-*his* (p-*abjA*), with and without the supplementation of 1 mM IPTG. (b) Affinity purification of MG1655 strain harboring pTrc-*abjA*-*his* and pCOLA-*trbE*-FLAG. Samples were subjected to SDS-PAGE stained with Coomassie brilliant blue (CB) and silver stain (SS). Specific bands corresponding to TrbE and AbjA were also excised and validated by MS, as well as immunoblot analyses using α-His and α-FLAG antibodies. (c) Cartoon representations of AlphaFold3-predicted TrbE (blue)-AbjA (red) model (ipTM = 0.74, pTM = 0.73), against a surface representation (transparency 50%) of hexameric TrbE revealing how AbjA could disrupt the interface of TrbE dimers. (d) Intracellular ATP concentration of various MG1655 strains expressing either or both TrbE and AbjA at mid-log as measured by the BacTiter-Glo(TM) Microbial Cell Viability Assay, without the supplementation of 1 mM IPTG on LB. Unpaired Student’s t-test: *, p <0.05, **, p < 0.01.

### AbjA directly interacts with and targets TrbE

Given the genetic data, we hypothesized that AbjA and TrbE physically interact, and furthermore that such an interaction results in cell death. Indeed, we found that TrbE-FLAG was stably co-purified with AbjA-His (Fig. 6b), establishing a direct interaction. AbjA is a protein of unknown function with no domain annotation. Within the T4SS, TrbE is a VirB4 homolog that forms a hexameric ATPase, with each protomer comprising an N-terminal globular and C-terminal ATPase domain, anchored to the membrane via a transmembrane helix. AlphaFold3 predicted an interaction model of AbjA and TrbE in a 1:1 ratio with high confidence; specifically, AbjA binds to the N-terminal globular domain of TrbE in a way that would disrupt oligomerization of the hexamer (Fig. 6c). Interestingly, intracellular ATP levels were also significantly reduced in strains co-expressing AbjA and TrbE (Fig. 6d). We therefore conclude that AbjA forms a complex with TrbE, possibly inducing formation of smaller oligomeric states with aberrant ATPase activity, eventually contributing to cell death by depleting intracellular ATP. This mode of action may explain why suppressor mutations in TrbE are largely truncations of the C-terminal ATPase domain.

## Discussion

Currently known defense systems that act against conjugation rely on nucleic acid recognition and degradation and are thus not specific to any aspect of the conjugation mechanism. AbjA thus represents the archetypal example of a new system with a novel mechanism-of-action, one which targets the T4SS apparatus directly. Since AbjA directly interacts with the ATPase TrbE to trigger cell death, and the latter is produced in the recipient cell only after plasmid transfer and establishment, AbjA appears to induce “abortive conjugation”. This mechanism is reminiscent of abortive infection against phage invasions, but has not been described for any non-phage plasmids. Like other immune systems, cell death induced by AbjA prevents subsequent plasmid transfer, thus limiting spread (infection) of the conjugative plasmid into the rest of the bacterial population. We have found that AbjA is present in a subset of *Enterobacteriaceae*, with distinct alleles in *Salmonella* that we suspect function by the same mechanism, suggesting that, similar to other defense systems like RM and CRISPR-Cas systems, there may be other naturally-occurring abortive conjugative factors that remain to be discovered.

It is intriguing that TrbE is the specific target of AbjA within the T4SS, as it represents the motor core of the conjugation machinery that drives pilus formation and/or DNA translocation in an ATP-dependent manner^42^. While widely variable components make up the T4SS across different families of conjugative plasmids, many of these are required for structure and function and thus would appear to be viable targets to disrupt the conjugation process. Our study demonstrates the feasibility of recipient-based factors targeting the next conjugation event to prevent plasmid spread. Furthermore, it reveals a possible additional advantage for targeting TrbE; beyond simply inhibiting its ATPase function, decoupling ATP hydrolysis from T4SS function can result in unregulated activity and abortive conjugation to effectively protect the bacterial population (Fig. 5 and 6). Accordingly, abortive conjugation may represent another previously unappreciated facet of the bacterial defense repertoire in the genetic ‘arms race’, which can be leveraged for the design of novel anti-conjugation therapeutics that might be useful for controlling the spread of AMR and virulence factors. Furthermore, because the conjugation machinery and other aspects of the conjugation process have not been seriously pursued pharmaceutically, they could represent a broad field of alternative drug targets, for a class of compounds we believe would complement existing antibiotic development and therefore term anti-resistics^43^.

## Methods

### Bacterial strains

All bacterial strains used in this study are listed in Supplementary Table S1.

### Plasmids and cloning procedures

Plasmids and primers used in this study are listed in Supplementary Table S2. Cloning procedures were performed following standard protocols of DNA digestion with restriction enzymes and ligation. All DNA constructs were sequenced by Sanger (1^st^ BASE, Singapore or Bio Basic Asia Pacific Pte Ltd, Singapore).

### Media and culture conditions

All strains were grown in Luria-Bertani broth (LB) or modified M9 medium (1× M9 salts, 1.5% Bacto agar, 2 mM MgSO_4_, 0.1 mM CaCl_2_, 0.2% rhamnose). Cultures were incubated at 37°C for 16-18 h (LB) or 40-42 h (M9). Antibiotics were added as needed to select for and maintain plasmids: ampicillin 200 μg/ml (Sigma-Aldrich, #A9518), kanamycin 50 μg/ml (#K1377), chloramphenicol 25 μg/ml (Calbiochem, #220551), tetracycline 10 μg/ml (#T7660), streptomycin 50 μg/ml (#S9137), spectinomycin 50 μg/ml (#S9007), and trimethoprim 10 μg/ml (#T7883). When using a rhamnose-inducible negative selectable marker, M9 medium was supplemented with 0.2% rhamnose (w/v) (#83650).

### Transformation assays

Electrocompetent cells were mixed with extracted conjugative plasmids and transferred to a 0.1 cm-gap Gene Pulser/MicroPulser electroporation cuvette (Bio-Rad, #1652089). Electroporation was performed using an Eppendorf Eporator (Eppendorf, #E4309000035) at 1,700 V. Each transformation contained 5 x 10^9^ to 1 × 10^10^ plasmid copies. Electroporated cells were recovered in LB supplemented with 2% glucose for 1 h, then plated on LB or modified M9 agar and incubated at 37°C for 16-18 h (LB) or 40-42 h (M9). Transformation frequency was calculated as the ratio of transformants over total recipient cells.

### Conjugation assays

Donor and recipient strains were grown overnight in LB supplemented with the appropriate antibiotics, shaking at 220 rpm and 37°C. Cultures were then subcultured 1:100 in fresh medium and grown to an OD_600_ of 1.0. Cells were pelleted and washed twice in LB without antibiotics, taking care to handle donor cultures gently to minimize pilus breakage. After the final wash, each pellet was resuspended in 100 μl LB and adjusted to an OD_600_ of 20-25. Conjugation was performed by mixing 80 μl donor with 20 μl recipient (D:R = 4:1) and spotting the mixture onto a pre-dried, pre-warmed LB agar plate in two 20 μl and six 10 μl spots. Plates were incubated at 37°C for 1 h to allow cells to conjugate. Spots were then scraped into 500 μl 1× PBS, gently mixed with an additional 500 μl 1× PBS, concentrated to 200 μl, serially diluted tenfold, and spotted onto selective LB or modified M9 agar. Plates were incubated at 37°C for 16-18 h (LB) or 40-42 h (M9). Conjugation frequency was calculated as the ratio of transconjugants over recipients.

### Efficiency of plating assays

Overnight bacterial cultures were washed twice in LB (without antibiotics) and adjusted to an OD_600_ of 1.0. Cultures were serially diluted in 1x PBS in a sterile 96-well PCR plate to generate seven tenfold dilutions. Aliquots of 2 μl or 10 μl from each dilution were spotted onto the indicated selective plates and incubated at 37°C for 16-18 h (LB) or 40-42 h (M9). Data shown are representative of at least two independent biological replicates.

### Growth curve assays

Overnight bacterial cultures were washed twice with 1x PBS and adjusted to an OD_600_ of 0.45-0.5. A 5 μl aliquot of culture (OD_600_ = 0.5) was mixed with 145 μl of the indicated medium in a well of a 96-well clear Costar plate (final starting OD_600_ ∼0.01; total volume 150 μl). Growth was monitored using a SpectraMax M5 plate reader (Molecular Devices, USA) in kinetic mode at 600 nm, with measurements taken every 20 min following a 5 s shake, over 24 or 48 h.

### Confocal microscopy

A stock solution of Hoechst 33342 dye (5 μg/ml; ThermoFisher, Invitrogen, #H21486) was mixed 1:1 with bacterial samples to yield a final concentration of 2.5 μg/ml. Samples were incubated at 37°C for at least 30 min and washed twice with 1x PBS. For microscopy, 1% agarose pads were freshly prepared and air-dried for at least 1 h. A 2-5 μl aliquot of the sample was then spotted onto the agarose pad. Imaging was performed on a Zeiss LSM710 confocal microscope using a 100× objective. All images shown are representative of at least two independent biological replicates.

### Affinity purification experiments

Briefly, for each strain, a 1.5 L culture was grown in LB broth at 37°C to mid-log. Cells were pelleted by centrifugation at 4700 x g for 20 min and then resuspended in 20-mL TBS containing 1 mM of PMSF, 100 µg/mL of lysozyme, and 50 µg/mL of DNase I. Cells were lysed with three rounds of sonication on ice (38% power, 1 s pulse on, 1 s pulse off for 3 min). Cell lysate was centrifuged at 24 000 x g for 1 h at at 4 °C to collect the membrane fraction, which was solubilized with an extraction buffer (20 mM Tris-HCl pH 8.0, 150 mM NaCl, 1%(w/v) n-dodecyl-β-D-maltoside (DDM, Calbiochem)) at 4 °C for 1 hr. Cell debris was removed by centrifugation at 25000 x g for 1 hr at 4 °C, and the solubilized membrane fraction was thereafter and loaded into a column packed with 1.5 mL of TALON cobalt resin (Clontech), pre-equilibrated with 20 mL of wash buffer (20 mM Tris-HCl pH 8.0, 150 mM NaCl, 0.05 %(w/v) DDM, 10 mM imidazole) in a column for 1 h at 4 °C with rocking. The mixture was allowed to drain by gravity before washing vigorously with 10 × 10 mL of wash buffer and eluted with 4 mL of elution buffer (20 mM Tris-HCl pH 8.0, 150 mM NaCl, 0.05 %(w/v) DDM, 200 mM imidazole). The eluate was concentrated in an Amicon Ultra 10 kDa cut-off ultra-filtration device (Merck Millipore) by centrifugation at 4000 x g to ∼100 μL. Concentrated eluate was mixed with equal amounts of 2× Laemmli reducing buffer, heated at 95 °C for 10 mins, and subjected to SDS-PAGE analyses and immunoblotting (described below). The resulting gel was stained with silver stain kit (SilverQuest™ Silver Staining Kit, ThermoFisher), and subsequently Coomassie Blue staining and visualized.

### Immunoblot analyses

SDS-PAGE gel was transferred onto polyvinylidene fluoride (PVDF) membranes (Immun-Blot® 0.2 μm, Bio-Rad) using the semi-dry electroblotting system (Trans-Blot® TurboTM Transfer System, Bio-Rad). Membranes were blocked using a casein blocking buffer (Sigma) diluted to 1x. Conjugated α-FLAG and α-His (to horseradish peroxidase) were acquired from Abcam. Luminata Forte Western HRP Substrate (Merck Milipore) was used to develop the membranes, and chemiluminescent signals were visualized by G:Box Chemi XT 4 (Genesys version 1.3.4.0, Syngene).

### ATP Quantification

In order to determine total ATP levels, each strain was cultured in 5 ml of LB at 37°C and 200 rpm till the exponential phase and normalized to OD_600_ = 0.5. Total intracellular ATP levels of the samples were measured using the BacTiter-Glo™ Microbial Cell Viability Assay Reagent (Promega, Madison, WI), based on the luciferase reaction. Total intracellular ATP levels were determined by measuring luminescence levels. Sample values were extrapolated to an adenosine 5′-triphosphate sodium salt (Promega) standard curve of tenfold dilutions of ATP from 1 μM to 10 pM. Then, 100 µL of each culture is transferred to a fresh tube on a 96-well. For each strain, 100 μl of the supernatant was mixed with 100 μl of BacTiter-Glo™ Cell Viability Assay Reagent in a Corning® 96-well white polystyrene plate (Merck), and incubated at room temperature for 30 min. Luminescence was measured in a SpectraMax M5 (Molecular Devices, USA). All quantifications were carried out in triplicate in three independent experiments.

## Acknowledgements

We thank Natalie Jing Ma for providing us with the annotated pRK24 plasmid map. We also thank Yunn Hwen Gan (National University of Singapore) and Fernando de la Cruz (University of Cantabria) for their generous gifts of various conjugative plasmids. This work was supported by the Singapore Ministry of Health National Medical Research Council under its Clinician Scientist Individual Research Grant (MOH-000530) (to S.L.C. and S.-S.C.).

## Author Contributions

L.A.O.Y. performed all genetic and HGT experiments; J.Y. performed all biochemical experiments; S.T. conducted bioinformatics analyses; S.C. and C.G.X.C conducted initial transposon mutant screening; L.A.O.Y., J.Y., S.L.C., and S.-S.C. analyzed the data and wrote the paper.

## Conflicting Interests

The authors declare no competing interests.

## Notes

### Competing Interest Statement

The authors have declared no competing interest.

