## Supplementary Information for "A recipient-based anti-conjugation factor triggers an abortive mechanism by targeting the Type IV secretion system"

[7]Current affiliation: self

*Correspondence to:

This PDF includes: Supplementary Figures S1-S9

Supplementary Tables 1-6

Supplementary Methods

Supplementary References

### Supplementary Figures

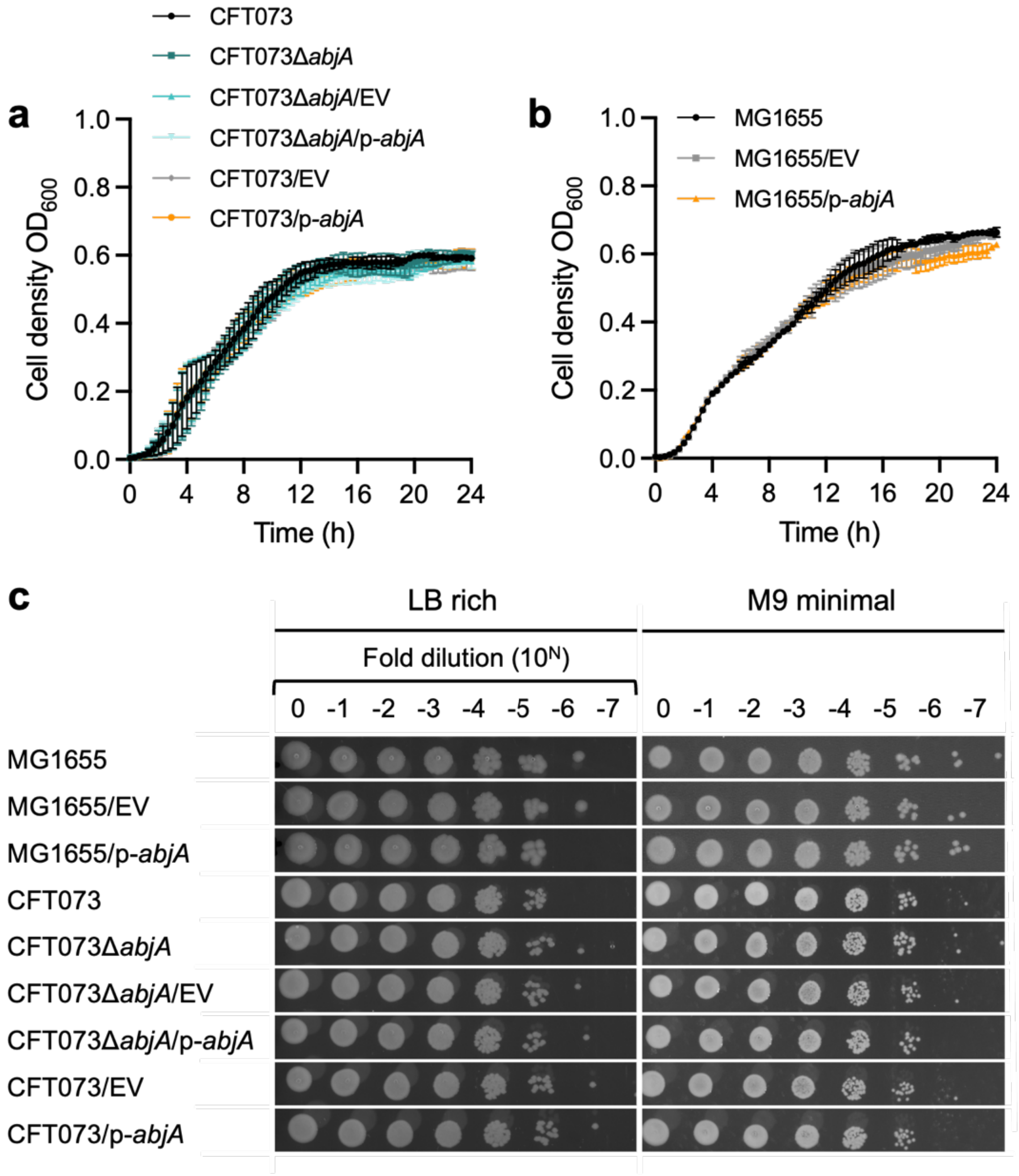


**Supplementary Figure S1. Presence of *abjA* does not impact fitness.** Growth curves of various (a) CFT073 and (b) MG1655 strains harboring pACYC (EV) or pACYC-*abjA*-*his* (p-*abjA*) in LB. (c) Efficiency of plating (EOP) of indicated strains in LB rich, or M9 minimal, media.

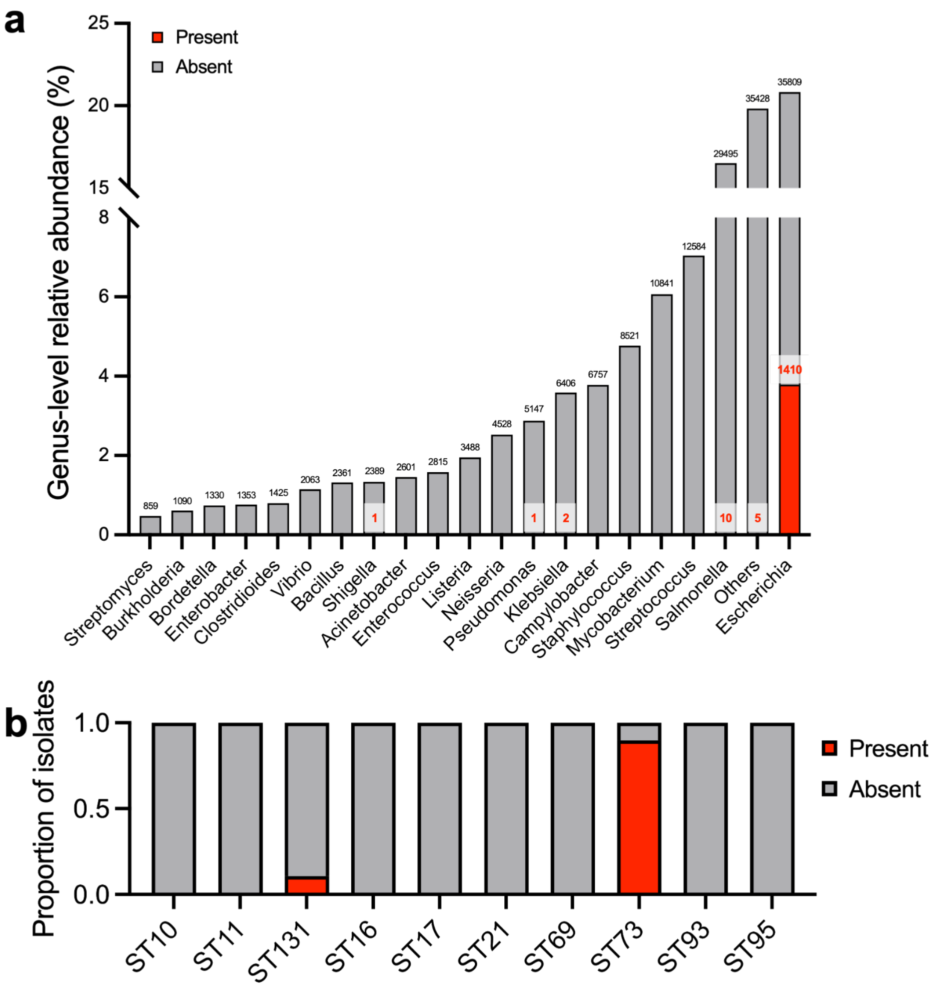


| ST | ST10 | | ST11 | | ST131 | | ST16 | | ST17 | | ST21 | | ST69 | | ST73 | | ST93 | | ST95 | |
|---|---|---|---|---|---|---|---|---|---|---|---|---|---|---|---|---|---|---|---|---|
| Dataset | Y | N | Y | N | Y | N | Y | N | Y | N | Y | N | Y | N | Y | N | Y | N | Y | N |
| LOGAN 122k | 8 | 1232 | 0 | 964 | 21 | 745 | 0 | 227 | 0 | 351 | 0 | 423 | 0 | 214 | 297 | 14 | 0 | 143 | 0 | 147 |
| NCBI SRA19k | 7 | 1475 | 0 | 2443 | 32 | 1600 | 0 | 371 | 0 | 601 | 0 | 1046 | 0 | 338 | 260 | 55 | 0 | 490 | 1 | 360 |
| NCBI RefSeq36k | 3 | 311 | 0 | 256 | 232 | 8 | 0 | 13 | 0 | 10 | 0 | 29 | 0 | 44 | 45 | 0 | 0 | 53 | 0 | 61 |
| ECOR72 | 0 | 9 | 0 | 0 | 0 | 0 | 0 | 0 | 0 | 0 | 0 | 0 | 0 | 1 | 6 | 0 | 0 | 0 | 0 | 0 |

**Supplementary Figure S2. *abjA* is largely limited to the *Escherichia* genus, and in sequence type 73 (ST73).** (a) Bar plot showing the top 20 bacterial genera identified based on taxonomy across the combined four datasets (LOGAN122k, NCBI RefSeq36k, NCBI SRA19k, ECOR72; see **Source Data** for details), where genera from datasets with full-length alignment to *abjA* (810 bp) are highlighted in red. Total numbers of genomes within each bacterial genera, and the number containing *abjA* are indicated at the top of the corresponding bars. (b) Bar plot illustrating the fractional distribution of *abjA* (highlighted orange) in the top 10 MLST sequence types (STs) for the Escherichia genus across the four datasets, as tabulated. Y = *abjA* present; N = *abjA* absent.

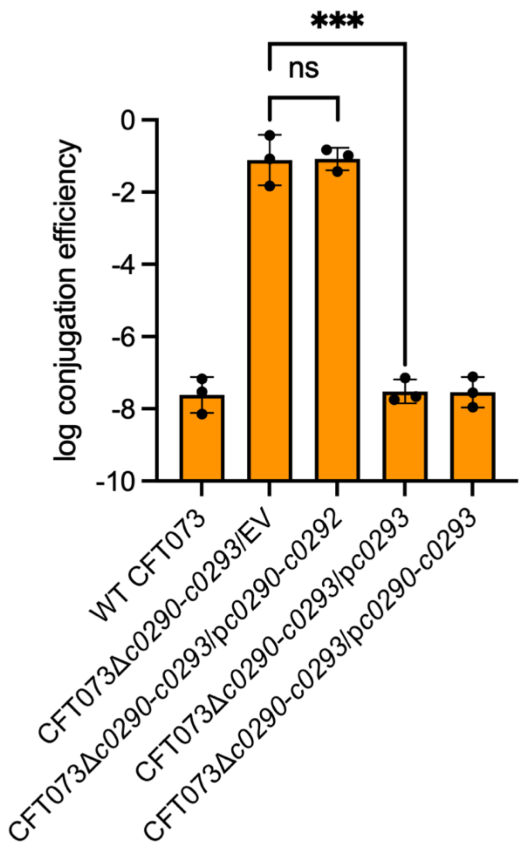


**Supplementary Figure S3. Adjacent genes *c0290–c0292* do not contribute to the conjugation inhibition phenotype.** Conjugation frequency of pRK24 into various recipient strains; all plasmids are based on a pACYC184 backbone. Log conjugation frequency data are presented as mean ± s.d. of three biological repeats. Unpaired Student's t-test: ***, $p < 0.001$, ns, not significant.

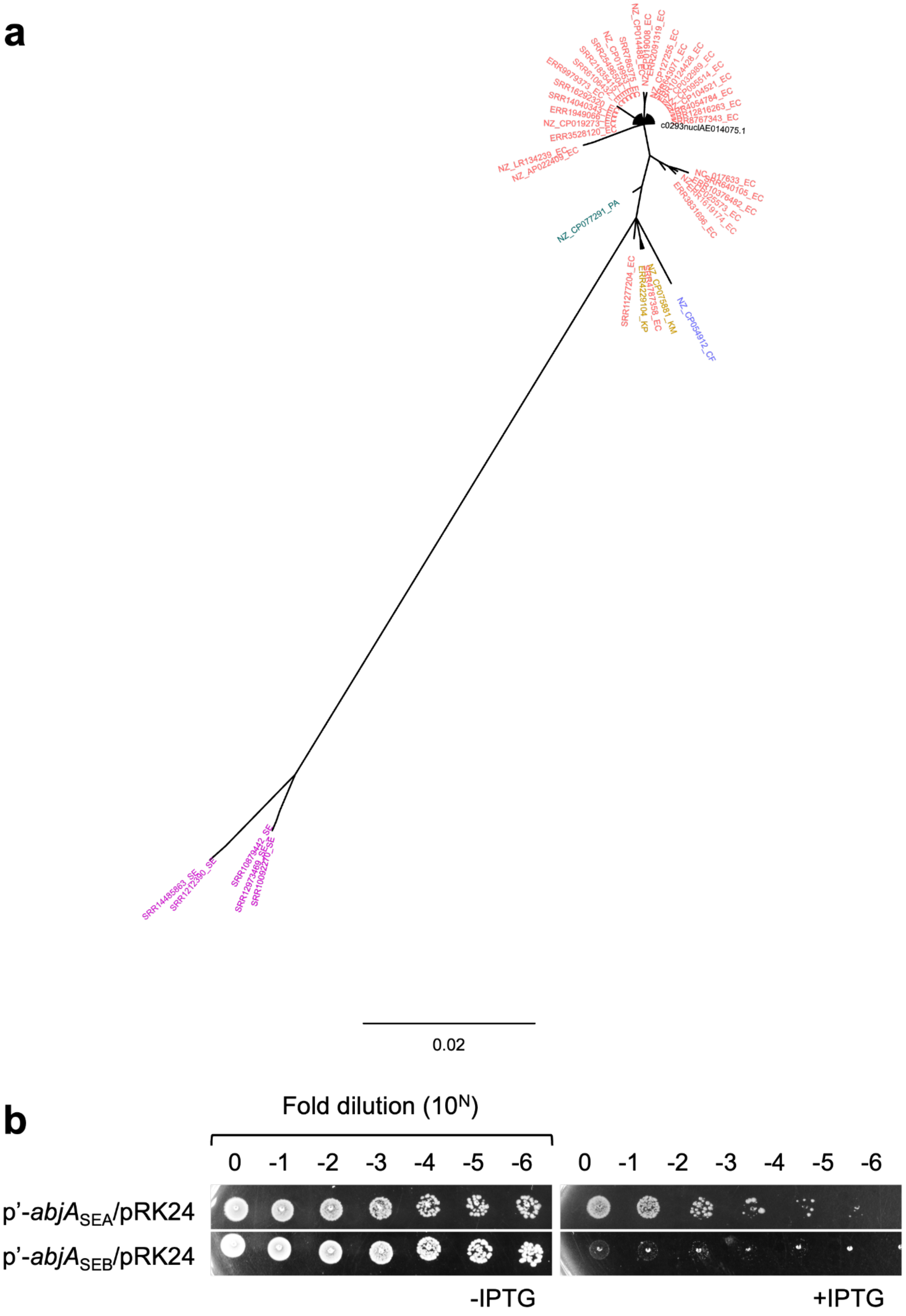
a
c0293nuclAE014075.1
NZ_CP077291_PA
SRR11277204_EC
NZ_CP054912_CF
0.02
b
Fold dilution (10^N)
0 -1 -2 -3 -4 -5 -6
0 -1 -2 -3 -4 -5 -6
p'-abjA_SEA/pRK24
p'-abjA_SEB/pRK24
-IPTG
+IPTG

**Supplementary Figure S4. *Salmonella* AbjA homologs are able to inhibit transfer of pRK24 into *E. coli.*** (a) Unrooted phylogenetic tree based on *abjA* single-gene nucleotide alignment, highlighting genus-level clustering among bacterial isolates. Tip labels indicate genome accession IDs, with suffixes (_EC, _CF, _KP, _KM, _PA, _SE) denoting the corresponding species. Genus codes: CF – *Citrobacter freundii*, EC – *Escherichia coli*, KM – *Klebsiella michiganensis*, KP – *Klebsiella pneumoniae*, PA – *Pantoea ananatis*, SE – *Salmonella enterica*. The scale bar (0.02) represents nucleotide substitutions per site. (b) EOP of MG1655 strains harboring *Salmonella abjA* allele A, pTrc-*abjA*$_{\mathrm{SEA}}$-*his* (p'-*abjA*$_{\mathrm{SEA}}$), and *Salmonella abjA* allele B, pTrc-*abjA*$_{\mathrm{SEB}}$-*his* (p'-*abjA*$_{\mathrm{SEB}}$), together with pRK24, with and without the supplementation of 1 mM IPTG on LB.

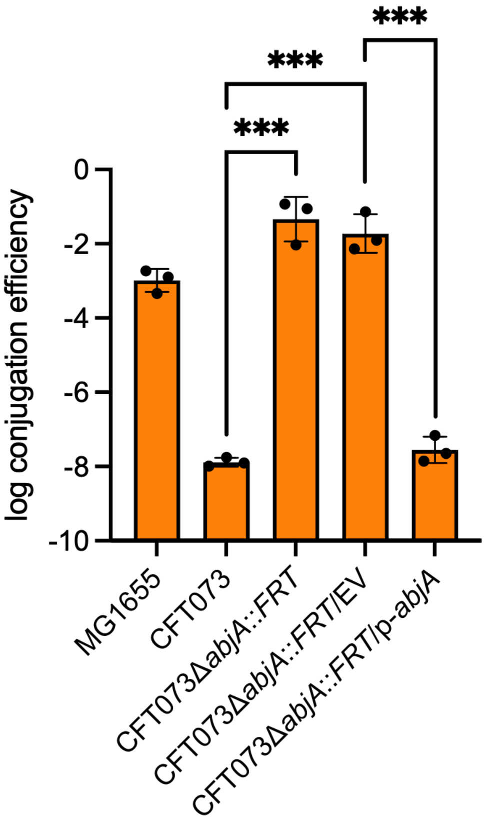


**Supplementary Figure S5. *abjA*-mediated inhibition of conjugation is independent of restriction-modification systems.** Conjugation frequency of pRK24, from a CFT073 donor strain, into MG1655, CFT073, CFT073Δ*abjA*, CFT073Δ*abjA*/pACYC177 (EV), and CFT073Δ*abjA*/pACYC177-*abjA*-*his* (p-*abjA*). Log conjugation frequency data are presented as mean ± s.d. of three biological repeats. Unpaired Student's t-test: ***, $p < 0.001$.

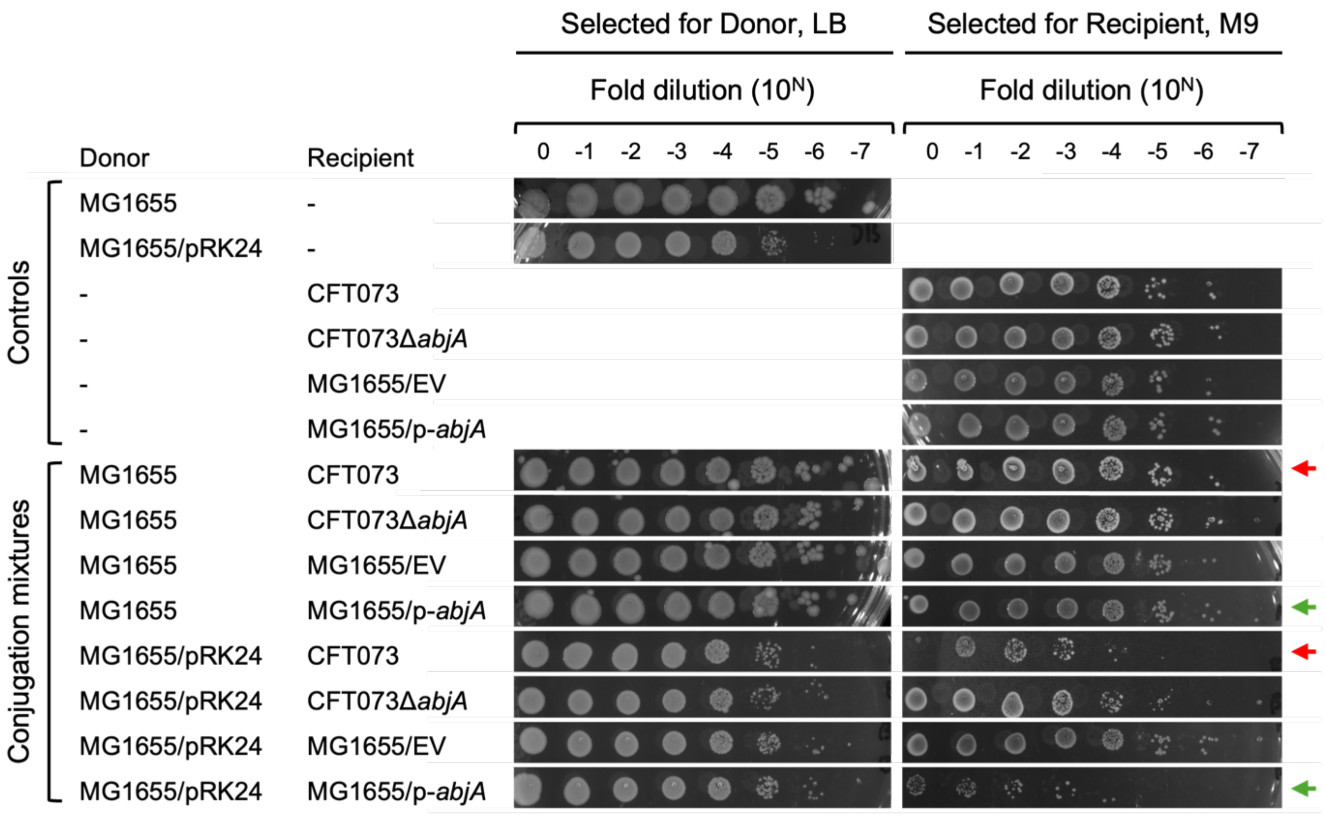


**Supplementary Figure S6. Co-occurrence of pRK24 and *abjA* reduced recipient CFU by 1-2 log.** EOP after conjugation is shown for donor-only controls, recipient-only controls, and donor-recipient mixtures of MG1655 or CFT073 carrying pRK24, pACYC177 (EV), or pACYC177-*abjA*-*his* (p-*abjA*). Selective plating used appropriate antibiotics and 0.2% rhamnose where required. Red and green arrow pairs indicate key ±pRK24 comparisons in CFT073 (*red*) and MG1655 (*green*).

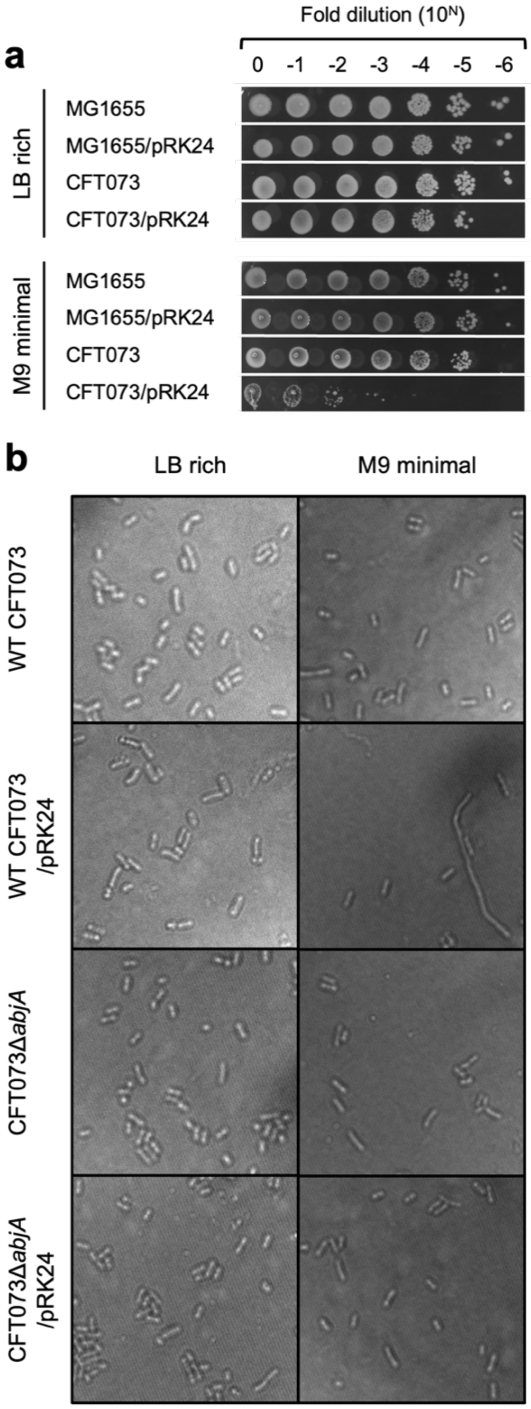


**Supplementary Figure S7. *abjA* phenotypes were observed in native host CFT073, but only in low nutrient conditions.** (a) Efficiency of plating of CFT073 and MG1655 carrying pRK24 on LB and minimal media agar plates. (b) Representative differential interference contrast (DIC) microscopy images of WT CFT073, WT CFT073/pRK24, CFT073Δ*abjA*, and CFT073Δ*abjA*/pRK24, visualized after being exposed to the standard conjugation conditions, LB (1 day incubation) or M9 (2 day incubation).

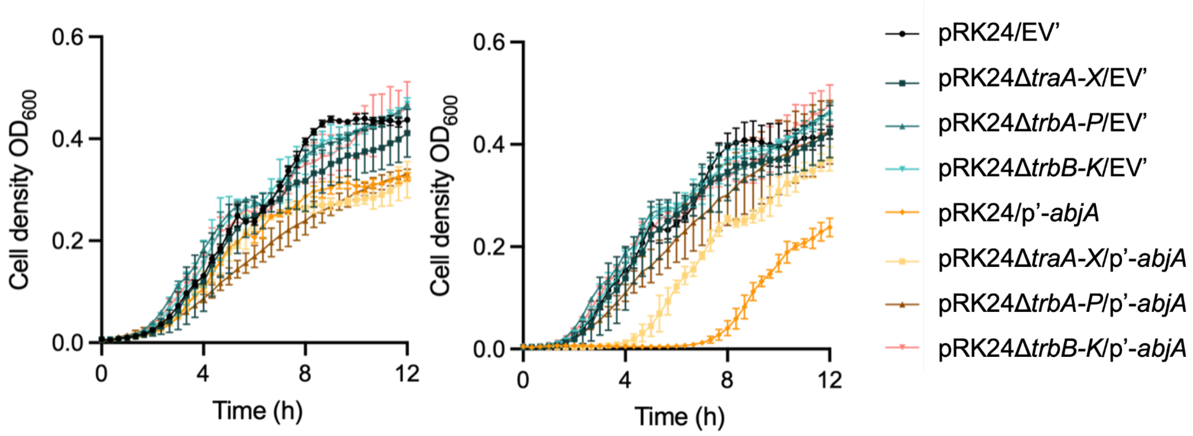


**Supplementary Figure S8. pRK24 Tra2 region is necessary for the lag growth phase.** Growth curves of MG1655 strains harboring pTrc99A empty vector (EV') or pTrc-*abjA-his* (p'-*abjA*), with indicated pRK24 deletion constructs, with and without the supplementation of 1 mM IPTG in LB.

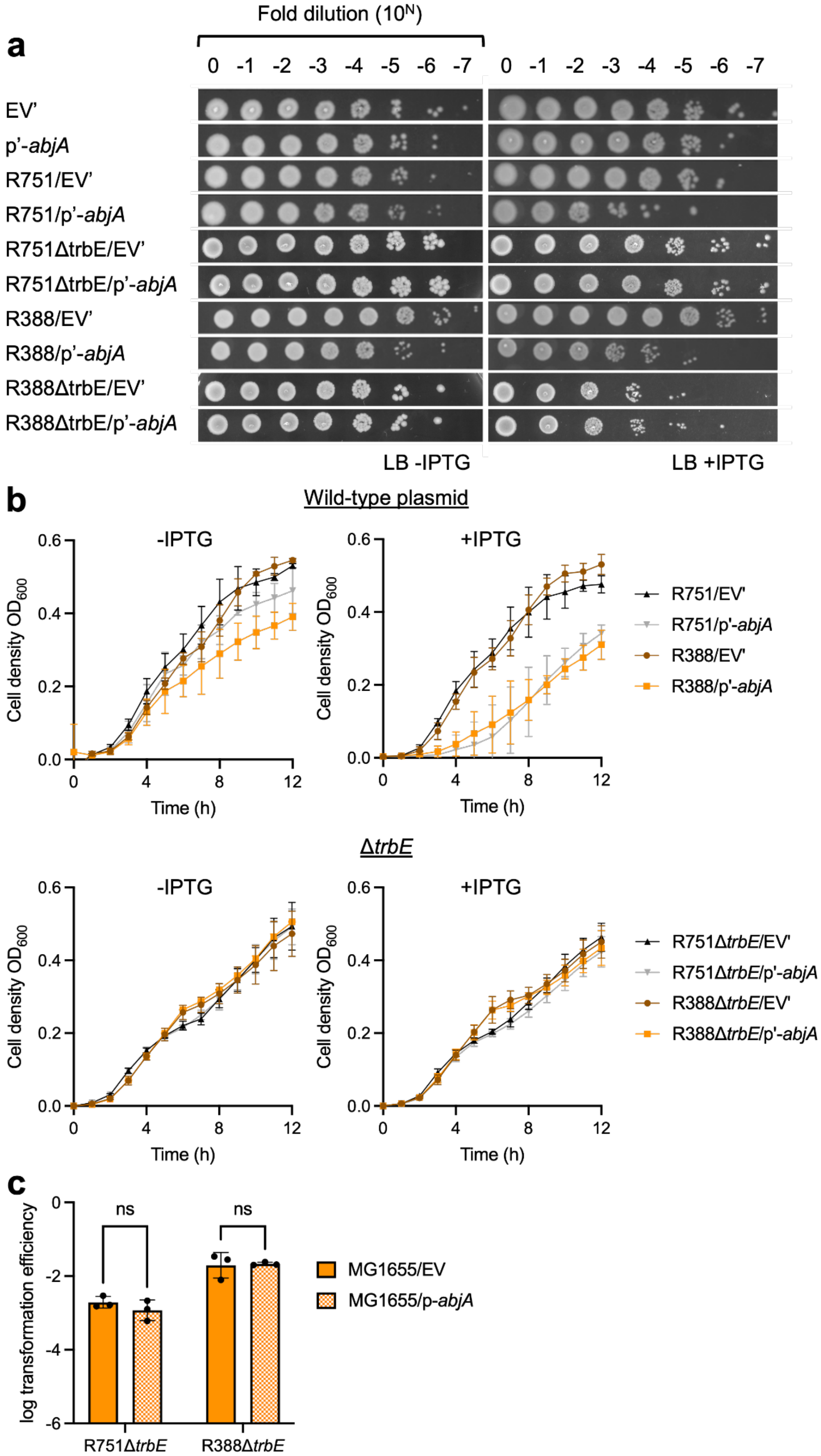
a
Fold dilution (10^N)
0 -1 -2 -3 -4 -5 -6 -7
EV'
p'-abjA
R751/EV'
R751/p'-abjA
R751ΔtrbE/EV'
R751ΔtrbE/p'-abjA
R388/EV'
R388/p'-abjA
R388ΔtrbE/EV'
R388ΔtrbE/p'-abjA
LB -IPTG
LB +IPTG
b
Wild-type plasmid
-IPTG
+IPTG
Cell density OD600
Time (h)
R751/EV'
R751/p'-abjA
R388/EV'
R388/p'-abjA
ΔtrbE
R751ΔtrbE/EV'
R751ΔtrbE/p'-abjA
R388ΔtrbE/EV'
R388ΔtrbE/p'-abjA
c
log transformation efficiency
ns
MG1655/EV
MG1655/p-abjA
R751ΔtrbE
R388ΔtrbE

83

**Supplementary Figure S9. TrbE is also a crucial factor on R751 and R388 that is singularly responsible for plasmid transfer inhibition.** (a) Efficiency of plating and (b) growth curves of MG1655/R751, MG1655/R751Δ*trbE*, MG1655/R388, and MG1655/R388Δ*trbE* harboring pTrc (EV') or pTrc-*abjA-his* (p'-*abjA*), with and without the addition of 1 mM IPTG on LB agar plates. (c) Transformation efficiency of R751Δ*trbE* and R388Δ*trbE* into MG1655 recipient harboring pACYC177 empty vector (EV) or pACYC177-*abjA-his* (p-*abjA*). Log conjugation frequency and transformation efficiency data are presented as the mean ± s.d. of three biological repeats. Unpaired Student's t-test: *, ns, not significant.

### Tables

**Supplementary Table S1.** Bacterial strains used in this study.

| Strain ID | Strain description | Reference/Source |
|---|---|---|
| BL21(λDE3) | *fhuA2 lon ompT gal* (λDE3) *dcm* Δ*hsdS* λDE3 = λ *sBamHIo* Δ*EcoRI-B int*::*(lacI*::*PlacUV5*::T7 *gene1*) *i21* Δ*nin5* | Novagen |
| NovaBlue | *endA1 hsdR17 (rK12– mK12+) supE44 thi-1 recA1 gyrA96 relA1 lac F' proA*$^{+}$ *proB*$^{+}$ *lacIq Z*Δ*M15*::*Tn10* | Novagen |
| SLC-1004 | MG1655Δ*HK*::*tse2-chlor* | [1] |
| SLC-1009 | CFT073Δ*HK*::*tse2-chlor* | [1] |
| SLC-6 | Wildtype UTI89 | [2] |
| SLC-7 | Wildtype MG1655 F$^{-}$ λ$^{-}$ *ilvG*$^{-}$ *rfb*-50 *rph*-1 | [3] |
| SLC-445 | Wildtype *Salmonella* Typhimurium 14028S | [4] |
| SLC-630 | Wildtype *Klebsiella pneumoniae* TOP1721 | [5] |
| SLC-948 | Wildtype CFT073 | [6] |
| SLC-1076 | CFT073 Tn5 C0293 clone1-13, pRK24 cured | This study |
| SLC-1077 | CFT073 Tn5 C0293 clone2-19, pRK24 cured | This study |
| SLC-1078 | CFT073 Tn5 C0293 clone3-14, pRK24 cured | This study |
| SLC-1156 | CFT073 Strep$^{R}$ | This study |
| LOY3E-30-7 | CFT073 Strep$^{R}$ Δ*abjA*::*FRT* | This study |
| SLC-1174 | CFT073 Strep$^{R}$ Δ*c0290-c0293*::*FRT* | This study |
| SLC-1155 | MG1655 Strep$^{R}$ | This study |
| SLC-1154 | UTI89 Strep$^{R}$ | This study |
| LOY3E-34-1 | *Salmonella* Typhimurium 14028S Strep$^{R}$ | This study |
| LOY3E-34-3 | *Klebsiella pneumoniae* TOP1721 Strep$^{R}$ | This study |
| LOY3A-40-1 | MG1655 Strep$^{R}$/pRK24-C1/pTrc99A | This study |
| LOY3A-40-3 | MG1655 Strep$^{R}$/pRK24-C2/pTrc99A | This study |
| LOY3A-40-7 | MG1655 Strep$^{R}$/pRK24-C3/pTrc99A | This study |
| LOY3A-40-2 | MG1655 Strep$^{R}$/pRK24-S1/pTrc-*abjA-his* | This study |
| LOY3A-40-4 | MG1655 Strep$^{R}$/pRK24-S2/pTrc-*abjA-his* | This study |
| LOY3A-40-8 | MG1655 Strep$^{R}$/pRK24-S3/pTrc-*abjA-his* | This study |
| LOY3A-40-9 | MG1655 Strep$^{R}$/pRK24-S4/pTrc-*abjA-his* | This study |
| LOY3A-40-10 | MG1655 Strep$^{R}$/pRK24-S5/pTrc-*abjA-his* | This study |
| LOY3A-40-11 | MG1655 Strep$^{R}$/pRK24-S6/pTrc-*abjA-his* | This study |
| LOY3A-40-12 | MG1655 Strep$^{R}$/pRK24-S7/pTrc-*abjA-his* | This study |
| LOY3A-40-13 | MG1655 Strep$^{R}$/pRK24-S8/pTrc-*abjA-his* | This study |
| LOY3A-40-14 | MG1655 Strep$^{R}$/pRK24-S9/pTrc-*abjA-his* | This study |
| LOY3A-40-15 | MG1655 Strep$^{R}$/pRK24-S10/pTrc-*abjA-his* | This study |

**Supplementary Table S2.** Plasmids used in this study.

| Plasmid ID | Plasmid name | Plasmid description | Reference/Source |
|---|---|---|---|
| EV | pACYC177 | Low copy cloning vector; p15A ori; Amp$^R$, Kan$^R$ | [7] |
| EV | pACYC177Δ*kan* | Low copy cloning vector; p15A ori; Amp$^R$ | This study |
| p-*abjA* | pACYC177-*abjA-his* | Low copy cloning vector; p15A ori; Amp$^R$<br>Used for expression of *abjA* under its native promoter; flanking intergenic regions were included, also where the promoter is assumed to be | This study |
| - | pACYC184 | Low copy cloning vector; p15A ori; Tet$^R$, Cam$^R$ | [7] |
| - | pACYC184-*c0290-c0292* | Low copy cloning vector; p15A ori; Cam$^R$<br>Used for expression of *c0290-c0292* under their native promoters; flanking intergenic regions were included, also where the promoters are assumed to be | This study |
| - | pACYC184-*c0290-c0293* | Low copy cloning vector; p15A ori; Cam$^R$<br>Used for expression of *c0290-c0293* under their native promoters; flanking intergenic regions were included, also where the promoters are assumed to be | This study |
| - | pACYC184-*c0293* | Low copy cloning vector; p15A ori; Cam$^R$<br>Used for expression of *abjA* under its native promoter; flanking intergenic regions were included, also where the promoter is assumed to be | This study |
| EV' | pTrc99A | High copy cloning vector; pBR322 ori; rrnB T2 terminator; Amp$^R$; for expression of genes under an IPTG-inducible pTrc promoter | [8] |
| p'-*abjA* | pTrc-*abjA-his* | High copy cloning vector; pBR322 ori; rrnB T2 terminator; Amp$^R$<br>Used for expression of *abjA* under an IPTG-inducible pTrc promoter | This study |
| p'-*abjA*$_{SEA}$ | pTrc- *abjA*$_{SEA}$-*his* | High copy cloning vector; pBR322 ori; rrnB T2 terminator; Amp$^R$<br>Used for expression of *Salmonella abjA* allele A (SRR10879442) under an IPTG-inducible pTrc promoter | This study |
| p'-*abjA*$_{SEB}$ | pTrc- *abjA*$_{SEB}$-*his* | High copy cloning vector; pBR322 ori; rrnB T2 terminator; Amp$^R$<br>Used for expression of *Salmonella abjA* allele B (SRR14485863) under an IPTG-inducible pTrc promoter | This study |
| EV'' | pCOLADuet-1 | Medium copy cloning vector; ColA ori; Kan$^R$ | Novagen |
| p''-*trbE*$_{pRK24}$ | pCOLA-*trbE*$_{pRK24}$ | Medium copy cloning vector; ColA ori; Kan$^R$<br>Used for expression of *trbE* under an IPTG-inducible T7 promoter | This study |

**Supplementary Table S3.** Plasmids used for the conjugation and transformation assays in this study.

| Name | Size (bp) | Inc group | Resistance | Isolated in | Genbank | Source |
|---|---|---|---|---|---|---|
| pRK24 | 60,099 | P1α | Tet, Amp | *Pseudomonas aeruginosa* and *Klebsiella aerogenes* | Sequence acquired from Ma and Isaacs (provided) | Chng lab |
| F plasmid | 99,159 | FI | Tet | *Escherichia coli* | NC_002483.1 | de la Cruz lab |
| drR27 | 180,461 | HI1 | Tet | *Salmonella enterica serovar* Typhimurium | NC_002305.1 | de la Cruz lab |
| R388 | 33,913 | W | Trim | *Escherichia coli* | NC_028464.1 | de la Cruz lab |
| R751 | 53,423 | P1β | Trim | *Enterobacter aerogenes* | U67194.4 | de la Cruz lab |
| pOX38 | 59,705 | FI | Cam | *Escherichia coli* | NZ_MF370216.1 | de la Cruz lab |
| pKPC2::*kan* | 71,861 | Unknown | Amp, Kan | *Klebsiella pneumoniae* | NZ_VONF01000004 | Gan lab |
| pCOLADuet-1 | 3,719 | ColA | Kan | - | Addgene #71406 | Chng lab |
| pBAD18-kan | 5,437 | pBR322 | Kan | - | Addgene #1841 | Chng lab |

**Supplementary Table S4.** Modifications made to conjugative plasmids investigated in this study. All modifications were made via the lambda Red recombineering method.

| Modified pRK24 | Gene(s) deleted | Gene(s) function category | Final antibiotic resistances |
|---|---|---|---|
| pRK24Δ*bla* (parent) | *bla* | Ampicillin resistance | Tet, Kan |
| pRK24Δ*traA-X* | *traA, traB, traC, traD, traE, traF, traG, traH, traI, traG, traH, traI, traJ, traK, traL, traM, traX, oriT, upf54-4 ORF, upf54-8 ORF* | DNA transfer processing | Tet, Kan, Spec |
| pRK24Δ*trbA-P* | *trbA, trbB, trbC, trbD, trbE, trbF, trbG, trbH, trbI, trbJ, trbK, trbL, trbM, trbN, trbP* | Mpf apparatus | Tet, Kan, Spec |
| pRK24Δ*klaABC* | *klaA, klaB, klaC* | Lethal genes | Tet, Kan, Spec |
| pRK24Δ*kleAB* | *kleA, kleB* | Lethal genes | Tet, Kan, Spec |
| pRK24Δ*kleCDEF* | *kleC, kleD, kleE, kleF* | Lethal genes | Tet, Kan, Spec |
| pRK24Δ*trbB-K* | *trbB, trbC, trbD, trbE, trbF, trbG, trbH, trbI, trbJ, trbK* | Mpf apparatus | Tet, Kan, Spec |
| pRK24Δ*trbK-P* | *trbK, trbL, trbM, trbN, trbP* | Mpf apparatus | Tet, Kan, Spec |
| pRK24Δ*trbBCD* | *trbB, trbC, trbD* | Mpf apparatus | Tet, Kan, Spec |
| pRK24Δ*trbEFG* | *trbE, trbF, trbG* | Mpf apparatus | Tet, Kan, Spec |
| pRK24Δ*trbHIJ* | *trbH, trbI, trbJ* | Mpf apparatus | Tet, Kan, Spec |
| pRK24Δ*trbE* | *trbE* | Mpf apparatus | Tet, Kan, Spec |
| pRK24Δ*trbF* | *trbF* | Mpf apparatus | Tet, Kan, Spec |
| pRK24Δ*trbG* | *trbG* | Mpf apparatus | Tet, Kan, Spec |
| R751Δ*trbE* | *trbE* | Mpf apparatus | Trim, Spec |
| R388Δ*trbE* | *HXF36_RS00035* | Mpf apparatus | Trim, Spec |

**Supplementary Table S5. Primers used in lambda Red recombineering.**

| Primer name | Sequence (5' to 3') | For deletion on |
|---|---|---|
| RK24 del(f7_bla) 5FP | TTACCAATGCTTAATCAGTGAGGCACCTATCTCAGCGATCTGTCTATTTCgtgtaggctggagctgcttc | pRK24Δf7 |
| RK24 del(f7_bla) 3RP | AATATTGAAAAAGGAAGAGTATGAGTATTCAACATTTCCGTGTCGCCCTTcatatgaatatcctccttag | |
| RK24 del(f6_full tra) 5FP | CTACCTCCGTAGTCGTAAGGTCGTTGCAGGTGCTCGGGTGCGGTACAACTtttttaaaacaatgaatagg | pRK24Δf6v2 |
| RK24 del(f6.2) RP | CCCGGCTCTAGGGAAAGCCGGCCTCGAACTGCCGAGGTCGGCTTTTTTTTttataattttttttaatctgt | |
| RK24 del(f5_spec) FP | CCGCTGTCAAGAACTTTAGCGGCTAAAATTTTGCGGGCCGCGACCAAAGGttttaaaacaatgaatagg | pRK24Δf5v2 |
| RK24 del(f5_spec) RP | CAGCAGAAAAAGCCGGGCATTGCCCGGCTTTCTTTGTGAGTCGTCCTGAAttataattttttttaatctgt | |
| RK24 del(f14_klaA-C) FP | ACGAGGGATAGAAAGTTTAGCTAAACTTCTTCCATCGAAAAGCAATTAACtttttaaaacaatgaatagg | pRK24Δf14 |
| RK24 del(f14_klaA-C) RP | TTGGCGTAACAAAACAAAAGCCCGGAAACCGGGCTTTCGTCTCTTGCCGCttataattttttttaatctgt | |
| RK24 del(f15_kleA-B) FP | CATAATGCCCTAATATAGCAATCCAAGGCCGGGCACTTCGCCCAGGTCAGttttaaaacaatgaatagg | pRK24Δf15 |
| RK24 del(f15_kleA-B) RP | GGGATTCAGCGAGGCGTCATGCTTGAAAACACCTTTCCCCTGGCGTGCAAttataattttttttaatctgt | |
| RK24 del(f16_kleC-F) FP | TAGGGCATTATGCCCTATTCTTGTTTTGAGGCCGGGTAGATTCCCAGGTCtttttaaaacaatgaatagg | pRK24Δf16 |
| RK24 del(f16_kleC-F) RP | ACACCTTTAGCCGCTAAAATTTGGGGACAGGTCATTTACAGAAAGCCAGCttataattttttttaatctgt | |
| RK24 del(f13_trbB-K) FP | ACACTTTCGGTATATCGTTTGCCTGTGCGATAATGTTGCTAATGATTTGTtttttaaaacaatgaatagg | pRK24Δf13 |
| RK24 del(f13_trbB-K) RP | TTTCGCTGGGCTTGAAGCCTCCCGGTTCTCCGCGCGGGCACTTCTTCAACttataattttttttaatctgt | |
| RK24 del(f2_spec) FP | CTGCGGGCGCGAAGCTGCGCGAGGGTTCGTACCGCGCAAGCCCGTCTAAGttttaaaacaatgaatagg | pRK24Δf2 |
| RK24 del(f2_spec) RP | GGGCCGCGATGGTGCATTTCTTGAGCAATTGCATATTGGAATCTCAAAGGttataattttttttaatctgt | |
| RK24 del(f18_trbBCD) FP | ACACTTTCGGTATATCGTTTGCCTGTGCGATAATGTTGCTAATGATTTGTtttttaaaacaatgaatagg | pRK24Δf18 |
| RK24 del(f18_trbBCD) RP | ACCGGCGGTGACGCAGGTACACGAACCGCATCTTCGGATCGGCCTTCGCCttataattttttttaatctgt | |
| RK24 del(f19_trbEFG) FP | TCCAAGCAATTGCGATTGCAATCGCGGGCCTCGGCGCGCTTCTGTTGTTCtttttaaaacaatgaatagg | pRK24Δf19 |
| RK24 del(f19_trbEFG) RP | TGGTCACGCGGTCCTGGCTGCTGCCCACGCCCGCGATGAGGATGGCCTTGttataattttttttaatctgt | |
| RK24 del(f20_trbHIJ) FP | ATGCGTAAGATTCTGACCGTCATCGCACTCGCGGCCACGTTGGCCGGCTGttttaaaacaatgaatagg | pRK24Δf20 |
| RK24 del(f20_trbHIJ) RP | CTTAGACGGGCTTGCGCGGTACGAACCCTCGCGCAGCTTCGCGCCCGCAGttataattttttttaatctgt | |
| RK24 del(f21_trbE) FP | TCCAAGCAATTGCGATTGCAATCGCGGGCCTCGGCGCGCTTCTGTTGTTCtttttaaaacaatgaatagg | pRK24Δf21 |
| RK24 del(f21_trbE) RP | ATCCACCCACTGGTCGCCGAACTTGGCTTCCAGGTTCTTGATGATGGCGAttataattttttttaatctgt | |
| RK24 del(f22_trbF) FP | GTTTTGCAGACACGATCAAGGGCTTGATCTTCAAGAAGAAGCCCGCAACGttttaaaacaatgaatagg | pRK24Δf22 |
| RK24 del(f22_trbF) RP | CGACGACGTAGACCGTCACCAAGGCCCGCATGCGCACGGGCTGGCCTTTCttataattttttttaatctgt | |
| RK24 del(f23_trbG) FP | ATGAAAAAGGAACTGTTTGCTTTGGTCCTGGCCGCGTCCGTTAGCGTGCCtttttaaaacaatgaatagg | pRK24Δf23 |
| RK24 del(f23_trbG) RP | TGGTCACGCGGTCCTGGCTGCTGCCCACGCCCGCGATGAGGATGGCCTTGttataattttttttaatctgt | |
| R751 del(f1_TrbE) FP | TCGAAGCAATCGCAATTGCCATTGCTGTGCTCGGTGCGGTTCTGTTGCTCtttttaaaacaatgaatagg | R751Δ*trbE* |
| R751 del(f1_TrbE) RP | ATGCACCCATTCGTGGCCGAACTTGGCTTCGAGGCTCTTGATGGTGGCGAttataattttttttaatctgt | |
| R388 del(f1_TrbERS035) FP | TGGGGCGCATCGTCGTATAGCCCCGCTAACTACCGAAAAAGGAGATAGGCtttttaaaacaatgaatagg | R388Δ*trbE* |
| R388 del(f1_TrbERS035) RP | CGCCGGGCTGTCTCCGACTTCGGCAATAATGCTTTCGGCAATTTCGGCATttataattttttttaatctgt | |

**Supplementary Table S6.** Mutations found in pRK24 and p'-*abjA* in the suppressor mutants.

| | **pRK24** | | | | **p'-*abjA*** |
|---|---|---|---|---|---|
| **Sample ID** | Plasmid ID | Mutation | Gene hit | Amino acid change | Mutation |
| **1** | pRK24-S1 | C21946del | *trbE* | p.G42AfsX14 | None found |
| **2** | pRK24-S2 | 22605C>T | *trbE* | Q261X | None found |
| **3** | pRK24-S3 | 23542G>A | *trbE* | W573X | None found |
| **4** | pRK24-S4 | A21988del | *trbE* | p.D55AfsX1 | None found |
| **5** | pRK24-S5 | 23664G>T | *trbE* | E614X | None found |
| **6** | pRK24-S6 | A21988del | *trbE* | p.D55AfsX1 | None found |
| **7** | pRK24-S7 | 23664G>T | *trbE* | E614X | None found |
| **8** | pRK24-S8 | A21988del | *trbE* | p.D55AfsX1 | None found |
| **9** | pRK24-S9 | 23542G>A | *trbE* | W573X | None found |
| **10** | pRK24-S10 | G22904del | *trbE* | A381X | None found |

### Supplementary Methods

#### Plasmid construction and cloning.

All plasmids used in this study are listed in Supplementary Table S2. To generate the plasmids, traditional ligation or Gibson assembly was used. In traditional ligation, the DNA inserts were amplified by PCR, then alongside the desired plasmid backbone, digested with the appropriate restriction enzymes (New England Biolabs). These were ligated with T4 DNA ligase (New England Biolabs, #M0202). In Gibson assembly, desired DNA fragments were inserted into linearized backbones using ClonExpress Ultra One Step Cloning Kit (Vazyme, #C115). In cases where the His-tag or FLAG-tag was inserted, primers used to amplify the DNA inserts were designed to also include a 6x His-tag (5'-CATCATCATCATCATCAT-3') sequence or 1x FLAG-tag (5'-DYKDDDDK-3') sequence downstream of the intended restriction cut sites/overlap regions.

#### Lambda Red recombineering.

Antibiotic resistance cassettes were amplified from pKD4 (kanamycin) or pSET2 (spectinomycin) using primers with ≥50 bp homology to the target chromosomal or plasmid insertion sites. Target cells harboring pKM208 (Addgene #13077) were subcultured 1:100 from overnight cultures, grown to $OD_{600}$ 0.26-0.30 at 30°C, 220 rpm, and induced with 1 mM IPTG to OD 0.4-0.5. Cells were heat-shocked at 42°C for 10-15 min with gentle agitation, cooled on ice for ≥10 min, and washed three times with ice-cold water to generate electrocompetent cells. Competent cells were electroporated with ≥1 µg DNA in 0.1 cm-gap cuvettes (Bio-Rad, #1652089) using an Eppendorf Eporator (Eppendorf, #E4309000035) at 1,500 V, recovered in LB with 2% glucose for 2 h at 37°C with shaking, followed by 2 h static, and plated on LB agar for 16-18 h. Colonies were screened on the appropriate antibiotic plates and verified by PCR. For Flp-*FRT*-mediated marker removal, recombinants were transformed with pCP20, incubated at 30°C for 1 h, and streaked on selective and non-selective plates (42°C to cure pCP20, 30°C to maintain) to confirm plasmid loss and marker removal by PCR.

**Large plasmid extraction.**

Conjugative plasmid DNA was isolated using a modified miniprep protocol. Reagents P1 (resuspension buffer), P2 (lysis buffer), and N3 (neutralization buffer) were obtained from the PureNA Biospin Plasmid Miniprep Kit (Research Instruments Pte Ltd, #KN01-250). Overnight bacterial cultures were centrifuged at 4,700 rpm for 10 min, and the pellet was resuspended in P1 buffer (5 ml culture: 300 µl buffer). Cells were lysed with 300 µl P2 buffer, neutralized with 350 µl P3 buffer, and centrifuged at 14,800 rpm for 10 min. The supernatant was transferred in 500 µl aliquots, mixed with 100 µl 3 M sodium acetate (pH 5) and 500 µl isopropanol, inverted 4-6 times, and incubated for ≥30 min to enhance yield. Samples were centrifuged at 4°C for 20 min at 14,800 rpm, and the resulting pellet was washed once with 1 ml 70% ethanol. After removal of ethanol and drying, the pellet was resuspended in nuclease-free water for storage.

**Qubit fluorometric quantification.**

For quantification of conjugative plasmids used in transformation assays, DNA concentrations were measured with a Qubit 2.0 Fluorometer (Invitrogen, #Q32866). A master mix was prepared by combining 198 µl buffer with 1 µl dye per sample (N × [198 µl buffer + 1 µl dye]). A standard curve was generated using the Qubit dsDNA High Sensitivity Assay Kit (Invitrogen, #Q32854) by mixing 190 µl master mix with 10 µl of each standard (0 ng/ml and 10 ng/ml). DNA samples were prepared by mixing 198 µl master mix with 2 µl sample and quantified against the standard curve.

**Preparation of electrocompetent cells for large plasmid transformation.**

Bacterial cultures were grown in LB medium with appropriate antibiotics to an $OD_{600}$ of 0.7-0.8 (higher cell densities are required for efficient transformation of large plasmids, as transformation efficiencies are lower). Cultures were then cooled on ice for ≥10 min and washed three times with ice-cold sterile water, centrifuging at 4,700 rpm for 10 min at 4°C each time. After the final wash, the cell pellet was resuspended in 1 ml cold 10% sterile glycerol (Sigma-Aldrich, #G5516) and aliquoted (50 ml culture concentrated into three 50 µl aliquots) for use in transformation assays. Competent cells were stored at -80°C if not used immediately.

**Bioinformatic analyses.**

***(A) Dataset Selection and Processing***

We utilized four distinct bacterial genome datasets: NCBI RefSeq36k, NCBI SRA19k, ECOR72, and LOGAN122k in this study (Source Data). Each dataset was curated and processed using specific criteria to ensure high-quality genomic data for analysis.

NCBI RefSeq36k Dataset: This dataset, downloaded from the NCBI Reference Sequence Database (RefSeq) on December 25, 2023, consists of 36,622 complete bacterial genomes. The smallest genome in the dataset belongs to *Candidatus Karelsulcia muelleri* (142,117 bp), while the largest is *Archangium violaceum SDU8* (13,213,980 bp).

NCBI SRA19k Dataset: This dataset comprises 19,962 *E. coli* genome assemblies, derived from 23,772 publicly available *E. coli* SRA runs (dated up to 2021). Genome assemblies were refined through quality filtering, prioritizing N50 values and complete genome lengths, and are part of an ongoing study (not yet published).

ECOR72 Dataset: A set of 72 *E. coli* genome assemblies, obtained from a previously published study (PMID: 30533715)

LOGAN122k Dataset: This dataset consists of 122,064 bacterial genome accession IDs, extracted from the *Logan Unitigs and Contigs of the Sequence Read Archive (SRA) on AWS accessed on 16 March 2025 from* https://registry.opendata.aws/pasteur-logan. The dataset was refined by leveraging metadata from the AWS S3 bucket (sra-pub-metadata-us-east-1/sra/metadata), which required no sign-in. The filtering criteria included bacterial genome taxonomic identifiers (taxids) to ensure exclusive selection of bacterial sequences.

***(B) Tools and Scripts***

To identify full-length *abjA* sequences (810 bp), standalone BLASTn was performed against available genome datasets. Multilocus sequence typing (MLST) was conducted using PubMLST (PMID: 30345391), utilizing BLASTn for housekeeping genes and phylogrouping by ClermonTyping (PMID: 29916797). For sequence clustering, standalone CD-HIT (PMID: 16731699) was used to group homologous *abjA* gene sequences based on identity thresholds. Multiple sequence alignment (MSA) was performed using MAFFT (PMID: 28968734), followed by maximum likelihood phylogenetic tree construction with IQ-TREE2 (PMID: 32011700), applying ModelFinder Plus (MFP) for optimal substitution model selection. Additional computational analyses were performed using custom Python and R scripts, tailored for alignment processing, data visualization, comprehensive analysis, structured data retrieval, and dataset-specific refinements to enhance genomic interpretations.
